# Global mapping of RNA-chromatin contacts reveals a proximity-dominated connectivity model for ncRNA-gene interactions

**DOI:** 10.1101/2022.09.02.506418

**Authors:** Charles Limouse, Owen K. Smith, David Jukam, Kelsey A. Fryer, William J. Greenleaf, Aaron F. Straight

## Abstract

Non-coding RNAs (ncRNAs) are transcribed throughout the genome and provide regulatory inputs to gene expression through their interaction with chromatin. Yet, the genomic targets and functions of most ncRNAs are unknown. Here we use chromatin-associated RNA sequencing (ChAR-seq) to map the global network of ncRNA interactions with chromatin in human embryonic stem cells, and the dynamic changes in interactions during differentiation into definitive endoderm. We uncover general principles governing the organization of the RNA-chromatin interactome, demonstrating that nearly all ncRNAs exclusively interact with genes in close three-dimensional proximity to their locus, and provide a model predicting the interactome. We uncover RNAs that interact with many loci across the genome, and unveil thousands of unannotated RNAs that dynamically interact with chromatin. By relating the dynamics of the interactome to changes in gene expression, we demonstrate that activation or repression of individual genes is unlikely to be controlled by a single ncRNA.

## Introduction

Cell identity is determined by the precise execution of lineage-specific gene expression programs^1^. These programs are controlled by coordinated signals from regulatory DNA sequences, transcription factors, histone modifications and variants, and 3D genome organization. The role of RNAs in modulating these programs is increasingly appreciated^2, 3^. Many classes of RNAs bind chromatin, collectively termed here chromatin-associated RNAs (caRNAs). These include long non-coding RNA(lncRNAs)^4, 5^, heterogeneous nuclear RNAs (hnRNAs)^6, 7^, enhancer-RNAs (eRNAs)^8–10^, transposable element (TE)-derived RNAs^11–14^, and other chromatin enriched RNAs (cheRNAs)^15, 16^. Yet, the function of these RNAs on chromatin remains largely unknown.

LncRNAs can orchestrate complex regulatory circuits, exemplified by XIST that acts as a core regulator of X-chromosome inactivation^17^, and KCNQ1OT1 that mediates allele-specific silencing of imprinted genes near its locus^18, 19^. In addition to lncRNAs, other classes of caRNAs have genome regulatory functions. For example, eRNAs can affect expression of neighboring genes through modulation of RNA polII elongation^20, 21^, or recruitment of transcriptional coregulators^22, 23^. Nascent pre-mRNAs can interact with chromatin binding proteins and locally regulate chromatin compaction^6, 24^, and TE-derived RNAs can silence immune response genes and hamper T-cell effector functions^25^. Furthermore, many proteins involved in controlling chromatin state^26–30^ and topology^23, 31^ have RNA-binding activity, suggesting additional roles for caRNAs in chromatin regulation. Despite these examples, which caRNAs have gene regulatory roles and their mechanisms of action remain to be determined^32^.

With the exception of a small number of caRNAs, we do not know the genomic loci where these RNAs act. As a result, we don’t understand the network of interactions between caRNAs and genes or its complexity. Transcription of both lncRNAs^33, 34^ and regulatory elements^9, 35–37^ exhibit strong tissue specificity such that the ncRNA-gene interaction network is also likely cell-state dependent, although this remains to be experimentally tested. Characterization of the network of human caRNA-gene interactions at the full transcriptome scale represents an important goal^25, 38–41^.

Here, we used chromatin-associated RNA sequencing (ChAR-seq) to map the RNA-chromatin interactome in H9 embryonic stem cells and definitive endoderm^42–44^. From these data we characterize the global architecture of this interactome, present a predictive model for most RNA-DNA chromatin interactions, and identify RNAs deviating from this model. We generate a detailed caRNA-gene interaction network that defines the set of caRNAs that interact with each gene based on physical proximity. These interactions encompass lncRNAs and many unannotated intergenic RNAs that may help prioritize specific caRNAs for future functional validation. Through analysis of the dynamics of the interactome during differentiation we find that regulation of gene expression by individual caRNAs is extremely rare.

## Results

To detect and map caRNA interactions with the genome, we performed ChAR-seq^42–44^, a proximity-ligation method that captures and sequences RNA-DNA contacts genome-wide (Fig. 1a). We performed ChAR-seq in human H9 embryonic stem cells (ES) before and after differentiation into definitive endoderm (DE) to understand how changes in the caRNA- chromatin interaction network might relate to activation or repression of cell-state specific genes. We validated our cell differentiation system by qPCR against cell-state marker genes and immunostaining, which revealed pure (>99%) ES and DE cell populations (Extended Data Fig. 1a,b)^45^.

**Figure 1.**
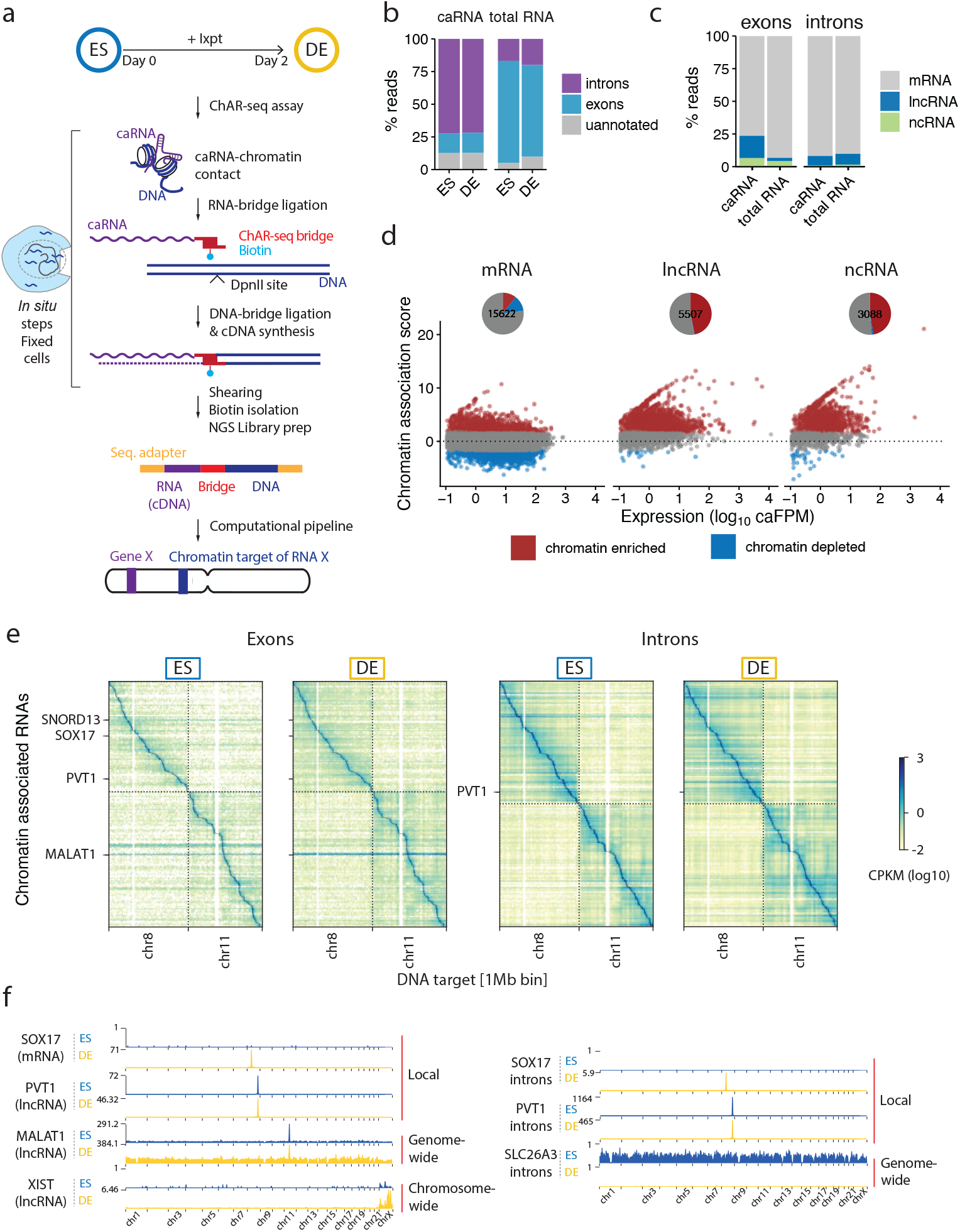
Global mapping of RNA-chromatin interactions during stem cell differentiation. a, Schematic of the strategy used to map RNA-DNA contacts across the transcriptome and genome using ChAR-seq, highlighting the key steps of the workflow. b-c, Composition of the caRNAs identified by ChAR-seq compared to the total RNA population determined by total RNA sequencing. d, Scatter plots showing the chromatin association scores for individual RNAs originating from annotated exons, as a function of the RNA level in the caRNA population. Chromatin enriched and depleted RNAs were determined using DESeq2 (FDR 0.05, fold change threshold 3x). Pie charts summarize the fraction of chromatin enriched and chromatin depleted RNA in each functional RNA type. The numbers within each pie chart indicate the total number of RNAs in that category. e, RNA-DNA contact maps in ES and DE cells for the top 200 most abundant caRNAs (according to their mean expression in ES and DE cells) on Chr7 and Chr8. Maps are displayed at a resolution of 1 RNA per row and 1 Mbp of genome space per column. Color represents contact density defined as the number of contacts between an RNA and a genomic bin, normalized for sequencing-depth and size of the genomic bin (CPKM: Contacts Per Kb in target genomic region per Million reads). Contacts made by exonic and intronic RNAs are shown in left and right maps, respectively. f, Interaction profiles along the genome for SOX17, PVT1, MALAT1 and XIST exons, and for SOX17, PVT1 and SLC26A3 introns, illustrating 3 major classes of interaction profiles: RNAs localized predominantly near their transcription locus (SOX17, PVT1 exons and introns), spreading across a single chromosome (XIST), and across the genome (MALAT1, SLC26A3 introns).

We sequenced ChAR-seq libraries to obtain over 900 million reads per cell state. We computationally split each read into a uniquely mapping RNA- and a DNA-derived sequence (Supplementary Note 1, Supplementary Fig. 1,2) and thereby obtained nearly 200 million unique RNA-DNA contacts (Extended Data Fig. 1c).

**Figure 2.**
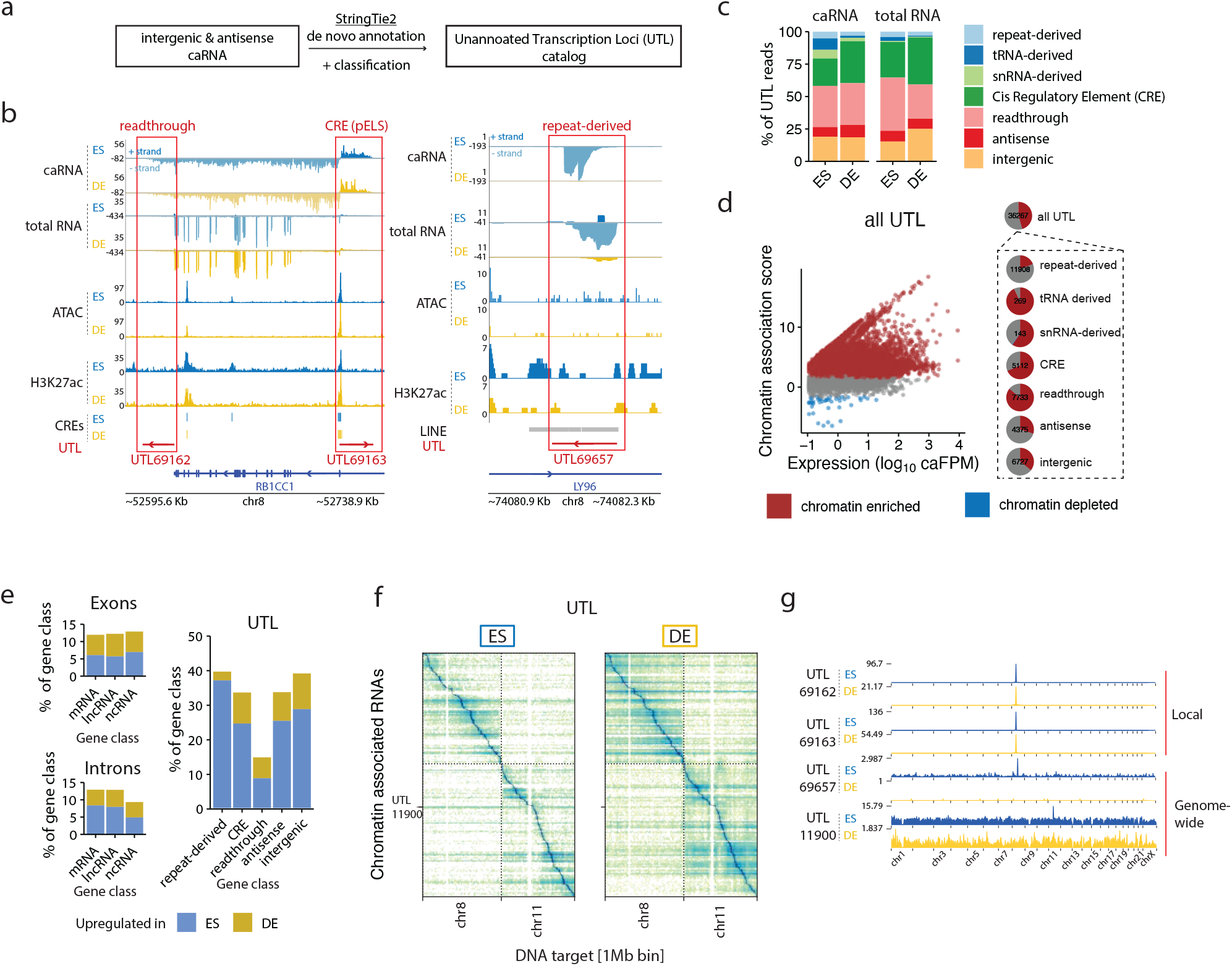
Cell state specific unannotated RNAs make up a large fraction of the caRNAs. a, Schematic of the method used to catalog unannotated RNAs by identifying transcription units using StringTie2. b, Genome tracks showing the chromatin context of 3 representative uannotated transcription loci (UTL). Left panel: UTL69162 and UTL69163, respectively downstream and antisense to RB1CC1 are classified as read-through RNA and CRE-derived RNAs. Right panel: UTL69657 is classified as a repeat-derived RNA due to its overlap with a LINE element. In both left and right panels, the top 2 tracks display the strand-specific genome coverage of the RNA-derived side of the ChAR-seq reads in ES and DE replicate 1 (+ strand ES in dark blue, - strand ES in light blue, + strand DE in dark yellow, - strand ES in light yellow). Next two tracks display the strand-specific genome coverage of the total RNA-seq data. c, Relative composition of the chromatin-associated UTLs in the 7 annotation classes. d, Scatter plots showing the chromatin association scores for individual UTLs and their abundance in the caRNA population. Chromatin enriched and depleted UTLs were determined using DESeq2 (FDR 0.05, fold change threshold 3x). Pie charts summarize the fraction of chromatin enriched and chromatin depleted UTLs in each category. Numbers within each pie chart indicate the total number of RNAs in that category. e, Percentage of genes upregulated and downregulated in DE vs ES cells in the caRNA transcriptome and for each RNA category. Up- and downregulated RNAs were identified using DESeq2 (FDR 0.05, fold change threshold 3x). f, RNA-DNA contact maps in ES and DE cells for the top 200 most abundant UTLs on Chr7 and Chr8, displayed at a resolution of 1 RNA per row and 1 Mbp of genome space per column. g, Genome-scale chromatin interaction profiles of 4 UTLs showing similar localization patterns as annotated RNAs.

We first analyzed the global composition of the caRNA population and found that caRNAs were enriched for non-coding RNAs, including introns, long non-coding RNAs (lncRNAs) and other functionally heterogeneous non-coding RNAs (referred to here as ncRNAs) such as small nuclear RNAs (snRNAs) and small nucleolar RNAs (snoRNAs; Fig. 1c, Extended Data Fig. 1d), consistent with previous studies^4, 46–48^. We normalized the caRNA population to expression levels by assigning each RNA a chromatin association score, defined as its relative abundance in the ChAR-seq versus total RNA-seq data (Methods). We found that nearly all introns and half of all non-coding RNAs had over 3-fold enrichment on chromatin, in agreement with prior characterizations of caRNA^16, 49^, indicating that ncRNAs tend to have nuclear or chromatin localization (Fig. 1d, Extended Data Fig. 1e, Table S2). LncRNAs are considered potential chromatin regulatory RNAs^3, 50^, yet our data indicate that non-intronic regions of lncRNAs constitute approximately 3% of the caRNA population, and less than 1% when excluding the top 10 most abundant lncRNAs. This result prompted us to perform a broad analysis of RNA-DNA interactions, including all caRNAs, rather than to focus exclusively on lncRNAs.

To compare the chromatin association patterns of exon- and intron-derived RNAs, we generated RNA-DNA contact maps for exons and introns (Fig. 1e). Our RNA-DNA contact maps were highly reproducible (Extended Data Fig. 1f) and showed high correlation between replicates and lower correlation between cell states, indicating that the interactome is dynamic during differentiation (Extended Data Fig. 1g). Across exons and introns, we uncovered several features of the RNA-DNA interactome mirroring those described in our prior work on *Drosophila melanogaster* and by others^43, 49, 51–53^. First, we noted a higher density of intrachromosomal compared to interchromosomal RNA-DNA contacts, reminiscent of the properties observed at the DNA level by Hi-C^54^, reflecting the chromatin organization into chromosome territories^55^.

Most RNA-DNA contacts occur close to the RNA transcription locus with on average ∼100-fold lower contact density 50-100 kb away from the transcription locus compared to at the transcription locus (Extended Data Fig. 1h). Finally, we observed three classes of RNA- chromatin association patterns (Fig. 1f). i) RNAs localizing predominantly at or near their transcription locus. ii) RNAs localizing across the genome, as previously observed^52, 56^. iii) RNAs such as XIST^57^ localizing across a single chromosome. We confirmed by RNA fluorescence in situ hybridization microscopy that the nuclear localization of select RNAs from these classes was consistent with their classification by ChAR-seq (Extended Data Fig. 2) and previous studies classifying noncoding RNAs by in situ hybridization ^58–62^. Altogether, these RNA- chromatin interactomes identify numerous RNAs in different functional classes that dynamically reorganize dependent upon cell state and demonstrate that most caRNAs remain associated with chromatin near their sites of synthesis.

### ChAR-seq identifies previously unannotated RNAs that bind chromatin dependent on cell state

We identified new unannotated RNAs that did not overlap with any known genes (as of Gencode v29) in 14% of all RNA-DNA contacts, a proportion similar to that of exons for annotated RNAs (Fig. 1b). To characterize the nature of these unannotated transcripts, we used the StringTie *de novo* transcriptome assembler to identify individual transcription units (Fig. 2a)^63^. We uncovered 30,442 loci with significant expression in ES or DE cells (FPM>0.1), which we hereafter refer to as unannotated transcribed loci (UTLs) (Extended Data Fig. 2b, Table S1, Table S3). Thus, the number of identified UTLs exceeds the number of known transcripts expressed at similar levels (22,475). We found that UTLs originated from functionally diverse chromatin loci (Fig. 2b). i) Some UTLs were immediately continuous with the 3’ end of active genes (e.g., UTL69162) and were possibly the result of transcriptional read-through, as reported in prior studies^64, 65^. ii) Some UTLs overlapped with regulatory signals such as high ATAC-seq or H3K27ac levels (e.g., UTL69163). iii) Some UTLs overlapped with TEs (e.g., UTL69657), in agreement with prior studies showing that TEs are a source of RNAs that associate with chromatin^11, 12, 25^. iv) Finally, some UTLs did not have any of the above features but had sequence similarity with known transfer RNAs (tRNAs), snRNAs and other small RNAs^66^.

Guided by these observations, we classified the UTLs based on their proximity to the 3’ or 5’ ends of genes, their overlap with transposable elements, snRNAs, or tRNAs, and their overlap with *cis* regulatory elements annotated in the Encode Registry of Regulatory Elements^67^, yielding 7 categories of unannotated RNAs (Methods, Table S3). ∼32% of the reads coming from UTLs were classified as readthrough RNAs and ∼27% as *cis* regulatory element-derived (Fig. 2c). Over 60% of the CRE-derived RNAs were from enhancer elements (Extended Data Fig. 2a). Four percent of the UTL reads were repeat-derived transcripts, roughly evenly distributed between LTR, SINE, and LINE elements (Fig. 2c, Extended Data Fig. 3a). Overall, the expression levels of UTLs were low, but similar to those of lncRNAs (Extended Data Fig. 3c).

Although these RNAs were present in the total RNA population, we found that all categories of UTLs were enriched on chromatin (Fig. 2d, Table S2) and were highly cell-state specific with 15- 49% of UTLs up- or down-regulated in the caRNA and total RNA populations compared to only ∼12% for mRNAs and lncRNAs (Fig. 2e). We examined the cell-state specificity and chromatin localization of two UTLs by fluorescence in situ hybridization and found that their localization was consistent with their ChAR-seq signal (Extended Data Fig. 3d). We generated RNA-DNA contact maps specifically for UTLs, which showed patterns similar to those observed for exonic and intronic RNAs (Fig. 2f). We found both UTLs which were locally restricted near their locus and UTLs that spread across the whole genome (Fig. 2g). This result prompted us to perform a broad analysis of all RNA-DNA interactions, including all caRNAs.

### RNA-DNA interactome dynamics is driven by caRNAs transcription dynamics rather than relocalization of caRNAs

We next quantified the dynamics of the RNA-chromatin interactome during ES-DE cell differentiation. To identify cell-state dependent interactions, we binned the DNA contacts of each RNA into 100 kb or 1 Mb intervals and performed a quantitative analysis analogous to differential expression analysis to obtain the fold change of each contact in ES versus DE cells and its associated statistical significance (Methods). We filtered the data to only include contacts with at least 10 counts in at least two samples, and tested ∼100,000 exon-chromatin contacts, ∼300,000 UTL-chromatin contacts, and 1.6 million intron-chromatin contacts (all at 100 kb resolution) for differential representation in ES vs DE cells (Extended Data Fig. 3a). The corresponding maps are shown in Fig. 3a. While we observed few dynamic RNA-chromatin interactions far from the RNA transcription locus (TL) in the exon and UTL maps, zooming in on a 10 Mb window around each RNA TL at 100 kb resolution revealed widespread changes in the interactome for all categories of RNAs. At 100 kb resolution ∼2% of interactions involving exons and ∼7% of interactions involving introns were up- or down-regulated in DE versus ES cells (Fig. 3b). More substantial changes were observed at a lower resolution of 1 Mb per genomic bin (Extended Data Fig. 3b). Consistent with the high cell state specificity of UTL expression discussed previously, UTLs also had the most dynamic RNA-DNA contact maps, with very low correlation between the ES and DE contact maps (Fig. 3b, Extended Data Fig. 4c).

**Figure 3.**
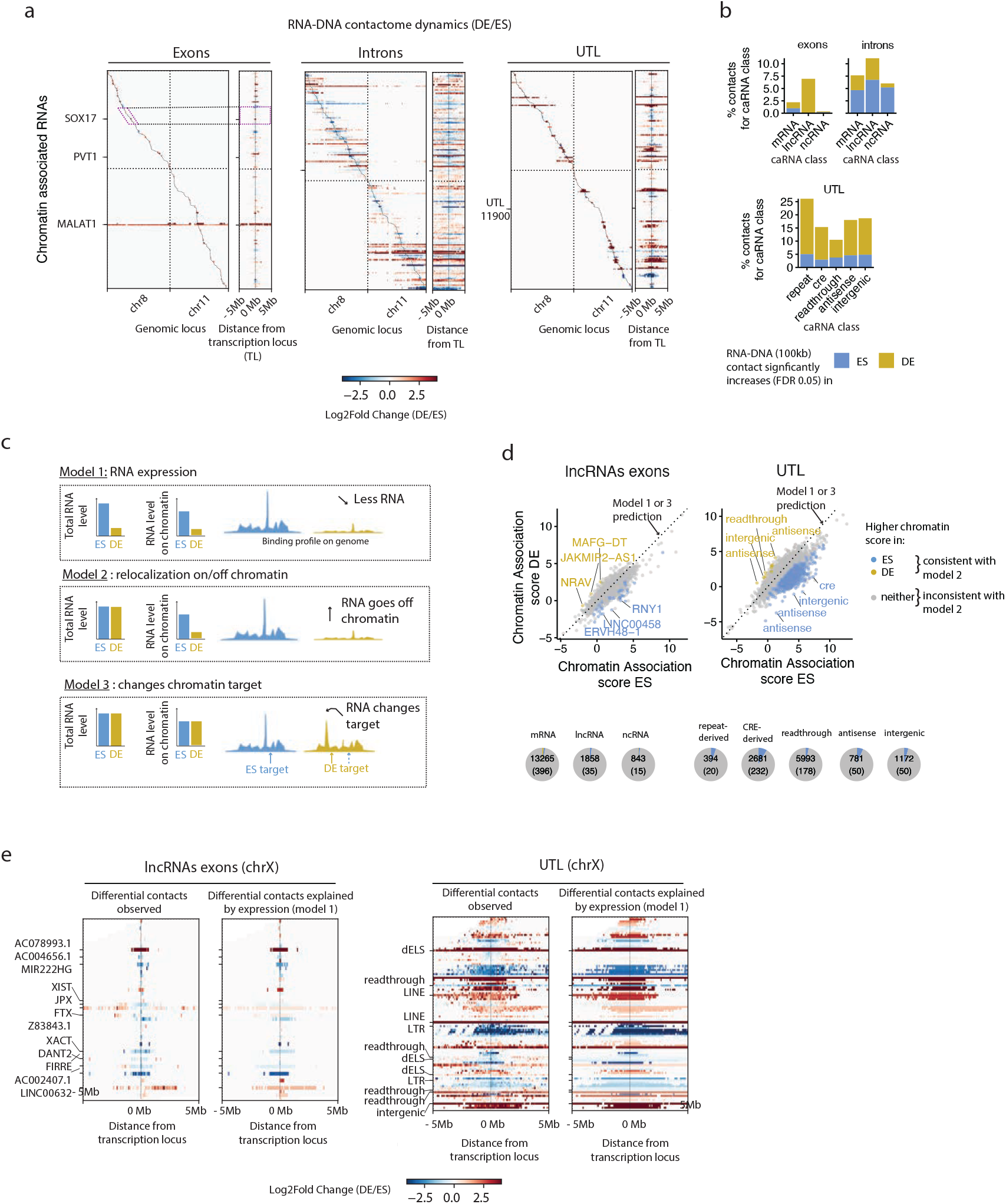
The RNA-DNA interactome dynamics are controlled at the transcription level. a, Differential contact maps showing the changes in the RNA-DNA interactome on Chr7 and Chr8 during cellular differentiation, for the same top 200 most abundant exonic RNAs, intronic RNAs, and UTLs as those shown Fig. 1e and Fig. 2f. For each RNA category, the left map shows the log2 fold change (LFC) in the frequency of each RNA-DNA contact, as computed by DESeq2 (shrunken LFC estimates, see Methods). x -axis resolution is 1 Mb as in Fig. 1e and Fig. 2f. The right map shows a zoom in of the left differential map in a 10 Mb window centered at the Transcription Locus (TL) of each caRNA, and displayed with an x -axis resolution of 100 kb. b, Quantification by RNA class of the percentage of interactions upregulated in DE or ES cells amongst all interactions tested in that class (interactions with >10 counts in at least one replicate in ES or DE), at 100 kb resolution (bottom panel). c, Schematic of 3 models that can explain changes in the DNA contact profile of an RNA during differentiation. d, Scatter plot showing the chromatin association score for individual lncRNAs exons (left panel) and UTLs (right panel) in ES versus DE cells. All of the caRNAs with an expression level above 0.1 FPM in both ES and DE cells are shown. Pie charts summarize the fraction of RNAs with significantly higher chromatin association in ES or DE cells (fold change >3, FDR 0.05), and for each RNA class. Numbers within the pie charts indicate the total number of RNAs in that class (FPM >0.1) and the number of RNAs with differential chromatin association. e, Differential contact maps observed versus those explained by transcription dynamics only for the 50 most abundant lncRNAs (left) and UTL (right) on ChrX. Labeled genes are the top 12 most abundant genes. x -axis resolution is 100 kb, and a 10 Mb window centered around each RNA TL is shown.

The interactome dynamics during differentiation may be driven by three non-mutually exclusive effects (Fig. 3c). First, an RNA may increase or decrease in overall abundance, resulting in proportionally increased or decreased binding levels on chromatin. Second, an RNA may modulate its affinity for chromatin, for instance, through RNA modifications or through changes in affinity with RNA-binding proteins mediating its interaction with chromatin. Third, an RNA may relocalize from one genomic site to another. The first two modes of dynamics would result in similar binding profiles in ES vs DE cells, albeit with an overall scale shift in binding levels. In contrast, the third mode implies changes in the RNA binding pattern to chromatin.

To test these models, we first compared the chromatin association score of each RNA in ES versus DE cells. Remarkably, the chromatin association scores remained mostly unchanged during differentiation, particularly for lncRNAs, with only 35 lncRNAs showing evidence of changes in their chromatin affinity (Fig. 3d, left panel, Table S2). Surprisingly, a larger fraction of UTLs, when compared to annotated non-coding RNAs (∼8% of CRE-derived UTLs and ∼5% of intergenic and antisense UTLs) showed significant changes in their chromatin association score between ES and DE cells (Fig. 3d, right panel). Thus, while individual RNAs show different propensities for chromatin interaction, this propensity does not change during differentiation and seems to be a property of the RNA itself. This result rules out model 2 for the majority of caRNAs.

Next, we examined whether the dynamics of specific interactions between an RNA and a chromatin locus can be explained by the transcriptional dynamics of the RNA itself. We compared the true differential contact maps to differential contact maps that would be observed if the frequency of each RNA-DNA contact was proportional to the total abundance of the corresponding RNAs in the caRNA population (Methods). These two differential interaction maps were highly similar (Fig. 3e). We further quantified the differences between these maps by identifying specific RNA-DNA contacts whose frequency changes between ES vs DE cells at a greater level than explained by the changes in RNA expression (Methods). We found no such contacts in the exon-DNA interactome and a negligible number of them in the UTL-DNA interactome (Extended Data Fig. 4d). Thus, the bulk of the changes in the RNA-DNA interactome appear to rely on transcription level regulation and expression differences in ES vs DE, rather than on modulation of an RNA’s affinity for chromatin or changes in an RNA’s contacts to different DNA binding sites.

### A select number of RNAs interact broadly with the genome

We hypothesized that the dynamic RNA-DNA interactome contains a mixture of i) functional interactions linked to regulatory activity of the RNA on chromatin, and ii) coincidental interactions due to transient proximity of the RNA to chromatin, for instance, during nascent transcription or diffusion within the nucleus. We thus analyzed the contact patterns of individual RNAs to detect features consistent with functional interaction, beginning with features at the chromosome scale. The nuclear speckle-associated lncRNA, MALAT1, and the XIST RNA are two well studied lncRNAs which act to regulate gene expression broadly across the genome or throughout the X chromosome^56, 62, 68^. Yet, it is not known which other RNAs have similar widespread interaction patterns on chromatin.

To systematically identify all RNAs with genome- or chromosome-wide associations, which we termed type I and type II RNAs (Fig. 4a), respectively, we developed two metrics, a *trans*- delocalization and a *cis*-delocalization score (Fig. 4b and Methods). The *trans*-delocalization score quantifies the tendency for an RNA to be found on chromosomes other than its source chromosome. Similarly, the *cis*-delocalization score assesses the tendency for an RNA to spread far (over 10Mb away) from its locus on its source chromosome. To account for expression, chromosome of origin and sample biases, these scores were calibrated using mRNAs as a reference (Methods, Supplementary Note 2, Supplementary Fig. 3]). We reasoned that type I RNAs must have high *trans*- and *cis*-delocalization scores, while type II RNA must have a high *cis*-delocalization score but a low *trans*-delocalization score. Thus, although other patterns may yield high delocalization scores (e.g. an RNA which targets a single locus on a *trans*-chromosome may have a large *trans*-delocalization score), we can use these metrics to screen for candidate RNAs with type I and type II patterns. We found that lncRNAs with large *trans*-delocalization scores (Fig. 4e, left panel) included MALAT1, the pTEFb-associated RNA, 7SK, and the telomerase RNA component, TERC, which all have established genome-wide chromatin regulatory functions, thus validating our approach^69–71^.

**Figure 4.**
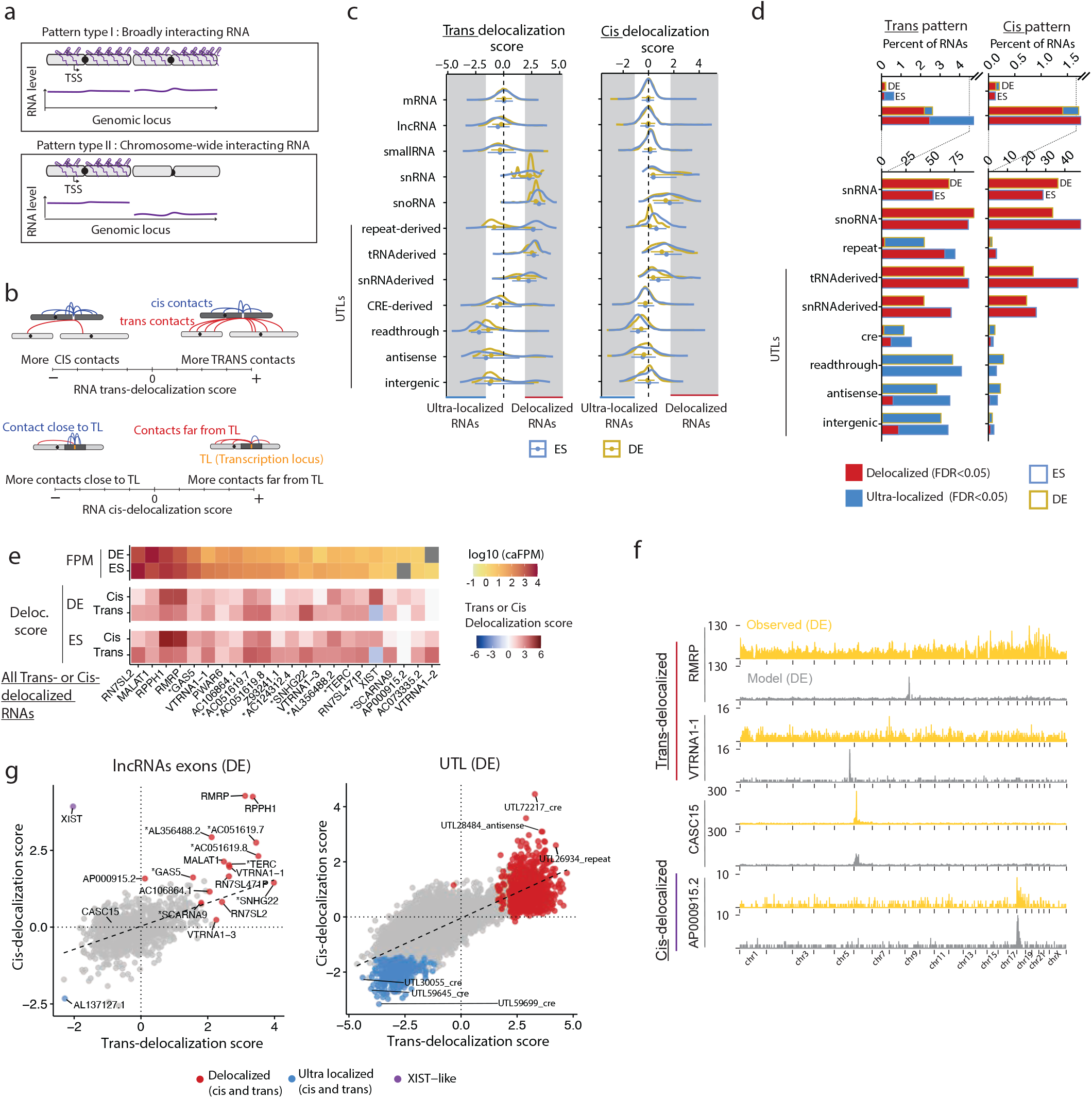
A select population of caRNAs interact with the genome broadly. a, Schematic of the two types of binding patterns identified in this analysis: type I) RNAs localized across the genome (trans-delocalized RNAs), type II) RNAs localized throughout their source chromosome but absent on other chromosomes (cis-delocalized RNAs). b, Schematic definition of the trans- and cis-delocalization scores. The trans-delocalization score quantifies the number of DNA contacts an RNA makes on chromosomes other than its source chromosome (trans contacts), relative to the number of contacts on its source chromosome (cis contacts). The cis-delocalization score quantifies the number of DNA contacts an RNA makes over 10 Mb away from its transcription locus (TL), relative to the number of contacts within 10 Mb of its TL. c, Distribution of trans- (left) and cis- (right) delocalization scores by class of RNA for exons and UTLs. d, Fraction of RNAs within each class identified as either delocalized or ultralocalized in regard to its trans- (left) or cis-chromosomal contacts (right). e, List of all lncRNAs identified as cis or trans-delocalized in either ES or DE cells and candidate RNAs for type I or type II patterns. Heat maps show the RNA cis and trans delocalization scores in ES and DE cells, and their abundance in the caRNA population. f, Chromatin interaction profiles for two examples of cis-delocalized RNAs (RMRP, VTRNA1-1), one example of cis-delocalized RNAs (AP000915.2), and one non-delocalized RNA (CASC15). Yellow track shows the observed ChAR-seq signal. Gray track shows the predicted interaction profile based on the generative model with trans-contact rate prediction, as described in Fig. 5 and Supplementary Note 4. g, Scatter plot showing the cis- versus trans-delocalization score for individual lncRNAs in ES cells (left), and UTLs in DE cells (right, excludes tRNA-derived and snRNA-derived UTLs). Colored data points indicate RNAs classified as delocalized (in either cis or trans), ultralocalized (in both cis and trans), and RNAs with XIST-like behavior. Black line shows the linear regression output.

We found that functionally distinct classes of RNAs had different distributions of delocalization scores (Fig. 4c, Table S4). LncRNAs had a wide range of delocalization scores, with a distribution of scores that mirrored those of mRNAs. In contrast, snRNAs, snoRNAs, tRNA- derived and snRNA-derived UTLs had globally high *cis*- and *trans*-delocalization scores, indicating that RNAs in these classes interact with loci throughout their source chromosome and across the whole genome. We observed the opposite behavior for CRE-derived RNAs and, to an even greater extent, for readthrough RNAs, which had mostly negative *cis*- and *trans*- delocalization scores, demonstrating that these RNAs tend to remain near their locus of origin. We also noted a negative-shifted distribution of delocalization scores for introns of both mRNAs and lncRNAs (Extended Data Fig. 5a). In ES cells, for ∼77% of individual lncRNAs and 96% of individual mRNAs, the *trans*-delocalization scores of their introns were lower than those of their exons (Extended Data Fig. 5b). Thus, introns tend to remain in closer proximity to their source locus.

**Figure 5.**
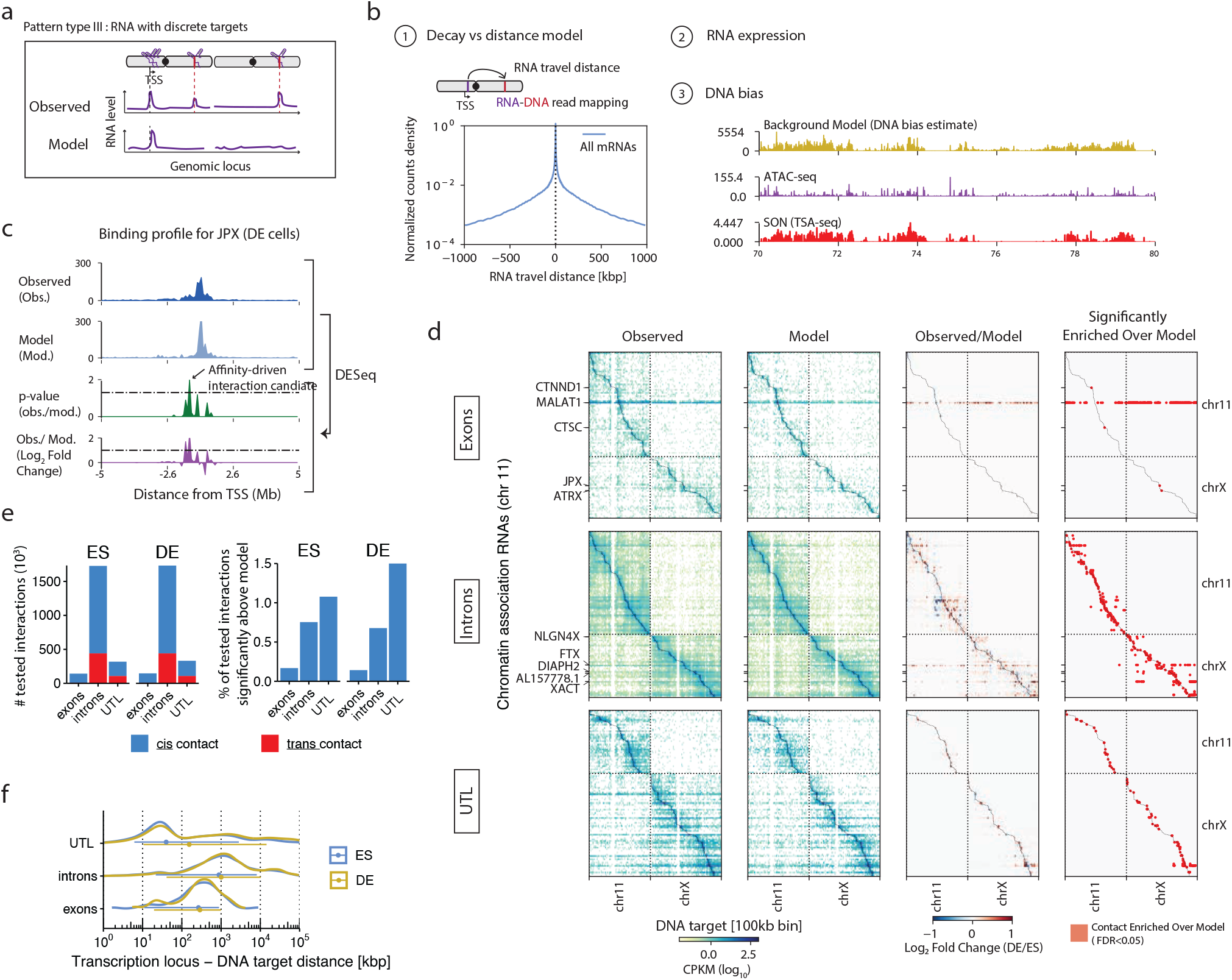
RNA expression and genomic distance determine the RNA-DNA interactome. a, Schematic of the type of binding patterns identified in this analysis. An RNA may localize at one or more discrete loci distinct from its transcription site (Pattern type III, top track) or remain in a diffusion constrained region around its locus (neutral RNA, bottom track). b, Components of the generative model used to predict the ChAR-seq maps. The number of contacts observed for an RNA at a DNA locus is proportion- al to i) an RNA-DNA distance-dependent contact frequency, ii) the abundance of the RNA on chromatin, iii) a target locus depen- dent bias (DNA-bias, yellow track) which correlates with both ATAC-seq signal (purple track) and nuclear speckle proximity signal (TSA-seq, red track). c, Example of a type III pattern with a candidate affinity-driven interaction for the lncRNA JPX in DE cells. The observed and predicted localization of JPX (top two tracks) at 10 kb resolution, and the are compared using DESeq2, yielding a Log2 fold change (observed over model) and an adjusted p-value track (bottom two tracks). Interactions with an LFC greater than 1.3 and an adjusted p-value smaller than 0.05 are labeled as “candidate affinity driven interaction”. d, Observed contact maps, predicted contact maps, and observed over model LFC maps computed using DESeq2 for the top 200 most abundant RNAs originating from exons (top), introns (middle) and UTLs (bottom). x -axis resolution is 100 kb per bin, Y-axis resolution is 1 RNA per bin. Only interactions with at least 10 counts in at least two samples were tested for differences with the model and are shown in the LFC maps. e, Number of interactions tested for enrichment over model and proportion of identified candidate affinity-driven interactions by RNA class, in relation to the total number of tested interactions in that RNA class. f, Distribution of the RNA-DNA travel distance for interactions significantly above model. The RNA-DNA travel distance is calculated using the mapping coordinates of the RNA and DNA side of the ChAR-seq read (Methods).

Interestingly, repeat-derived RNAs had globally high *cis*- and *trans*-delocalization scores in ES cells and low *cis*- and *trans*-delocalization scores in DE cells (Fig. 4c). Thus, in ES cells specifically, many repeat-derived RNAs tend to localize away from their transcription locus. To identify RNAs with extreme association scores, we applied an empirical Bayes method using mRNAs as a training set, which essentially identified RNAs in the 5% right-tail or the 5% left-tail of the mRNA score distribution (Method, Supplementary Note 3). We thus created a complete catalog of RNAs with candidate chromosome- or genome-wide association patterns, and another catalog of RNAs that remain localized within a 10 Mb window around their transcription locus or on their own chromosome, which we termed ultra-localized RNAs (from a *cis*- or *trans*- chromosomal perspective, Table S5). As expected, >50% of snRNAs, snoRNAs, tRNAs,and snRNAs were classified as *trans*-delocalized and >70% of readthrough RNAs were classified as ultra-localized (Fig. 4d). Surprisingly, out of 1,289 ncRNAs above 1 FPM with sufficient signal to compute delocalization scores (Methods), we detected only 22 lncRNAs (1.7%) with *cis*- or *trans*-delocalized patterns in either ES or DE cells (Fig. 4d, Extended Data Fig. 5c). In contrast, we found (excluding tRNA-derived and snRNA-derived UTLs) 60 UTLs in DE cells and 836 UTLs in ES cells and with *cis*- or *trans*-delocalization patterns, including 349 repeat-derived RNAs, and several hundreds of intergenic or CRE-derived UTLs (Extended Data Fig. 5c). The lncRNAs we characterized contained the known broadly acting RNAs discussed above.

Importantly, we discovered new candidate lncRNAs with potential genome-wide regulatory functions, including the mitochondrial RNA processing endoribonuclease RNA, RMRP, which is implicated in rRNA maturation^41, 72, 73^, the Ribonuclease P RNA Component H1, RPPH1, which is involved in tRNA processing^74, 75^, two isoforms of the Vault RNA, VTRNA1-1 and VTRNA1-3, and a large number of UTLs. We validated the delocalization score analysis by directly examining the ChAR-seq signal of these RNAs, which revealed their association across the genome (Fig. 4f). The delocalization of these RNAs was not explained by their abundance.

Although MALAT1, 7SK, and RMRP were highly abundant, other delocalized RNAs were all below 10 FPM. Furthermore, many abundant ncRNAs had low delocalization scores (Extended Data Fig. 5d). To confirm that the broad patterns detected by our delocalization score approach were not random or due to non-specific interactions, we performed metagene analysis centered on select genomic features. We detected enrichment of snRNAs at RNAPII occupancy loci (Extended Data Fig. 5e), where MALAT1 and 7SK were also enriched, consistent with the role of these RNAs in cotranscriptional splicing and transcriptional elongation^62, 69^. In contrast, VTRNA1-1 was found at background levels at RNAPII occupied loci, and RMRP was depleted at these loci. Together, our data show that broadly localized RNAs are rare amongst annotated lncRNAs, but we discovered a large repertoire of UTLs with potential global chromatin regulatory roles, specifically in ES cells.

While our characterized RNAs that were identified as significantly delocalized in *cis* but not in *trans*, we noted that amongst these RNAs, all but XIST also had a high *trans*-delocalization scores, albeit below the FDR threshold for classification as *trans*-delocalized. Generally, across all RNAs, the *cis*- and *trans*-delocalization scores were strongly correlated, indicating that RNAs that localize broadly on their own chromosomes also interact broadly with the rest of the genome (Fig. 4g). Remarkably, XIST was the only exception to this rule and was the only RNA which was simultaneously delocalized in *cis* and ultra localized in *trans*, consistent with its known localization throughout its source chromosome X (Fig. 4g). We concluded that XIST is unique in these cell types in its ability to interact with an entire chromosome while being excluded from other chromosomes.

We next examined changes in RNA delocalization in different cell states. We found that the delocalization scores were highly correlated between ES and DE cells, even for RNAs that were differentially abundant across cell states (Extended Data Fig. 4=5f). We thus concluded that the extent to which an RNA interacts with chromatin far from its transcription locus or on *trans* chromosomes is encoded in the RNA itself or the position of its transcription locus relative to other genomic features, rather than post-transcriptionally regulated.

### RNA-DNA contacts occur in the vicinity of the transcription locus

Engrietz et al. proposed a dichotomization of RNA-chromatin interactions into proximity-driven and affinity-driven interactions^2^. The former describes interactions occurring in a 2D or 3D distance bounded region around the transcription locus, without specificity for particular loci within that region. The latter describes RNA targeting well-defined loci, irrespective of their distance to the RNA locus. Some ncRNAs have been proposed to have affinity-driven interactions and regulate transcription or 3D organization of chromatin at their target loci^3, 76–78^. These data motivated us to search the interactome for contact patterns in which an RNA shows discrete peaks in its localization profile that are not explained by proximity to its locus (Fig. 5a, top panel, hereafter referred to as Type III patterns). Because standard genomic peak finding tools like MACS2^79^ are not appropriate for ChAR-seq data, we instead developed a generative model, which predicts the RNA-DNA interactome based on 3 features: 1) the total abundance of each RNA on chromatin, 2) a DNA-locus bias which models the propensity for an RNA to be captured at this locus, independently of the identity of that RNA, and 3) the distance between each RNA transcription site and its DNA target loci (Fig. 5b, Methods and Supplementary Note 4). As anticipated, the DNA-locus bias correlated with ATAC-seq, likely due to a combination of biological factors such as fewer RNA-DNA interactions existing in compact chromatin, and technical biases related to accessibility of the ChAR-seq bridge molecule. The DNA-locus bias also correlated with nuclear speckle proximity as measured by TSA-seq^80^, revealing a possible increased affinity for diffusing RNAs towards nuclear speckles. We trained our generative model on mRNAs, as we reasoned that most mRNAs should not have defined chromatin targets. We then used our final model to generate a “predicted” contact pattern for each RNA, which effectively provides a null hypothesis representing “neutral” patterns, where an RNA interacts exclusively and non-specifically with neighboring loci due to diffusion (Fig. 5a, model track).

Thus, positive deviations from the prediction (more contacts in the observed data compared with the model prediction) provide evidence for peak-like interactions in type III patterns.

In both ES and DE cells and for exons, introns, and UTLs, our simple generative model produced RNA-DNA contact maps highly similar to experimentally generated ChAR-seq RNA-DNA contacts maps (Fig. 5d, Extended Data Fig. 6a). At 100 kb DNA locus resolution and excluding RNAs previously identified as *cis*- or *trans*-delocalized, we identified only ∼0.2% of exon and ∼0.7% of intron contacts that were not explained by the model, irrespective of whether the RNAs were mRNAs, lncRNAs, or ncRNAs (Fig. 5e and Extended Data Fig. 6b-c). We detected only 11 and 9 lncRNAs in ES and DE cells, respectively, with exons making contacts in the genome at loci not predicted by our model (Table S6). Our model also accurately predicted changes in contact rates during differentiation (Extended Data Fig. 6d). Thus, in contrast with prior studies^76–78^, we found no evidence for type III patterns, where individual RNAs target distinct loci away from their transcription site, amongst the entire lncRNA population.

Interestingly, in contrast with that of lncRNAs, the interactome of the UTLs differed more substantially from its prediction. Over 1% of contacts involving 2,283 distinct RNAs in ES cells and 2,597 in DE cells showed statistical evidence for affinity-driven interactions (Fig. 5e). Readthrough RNAs had the largest number of such contacts followed by CRE-derived RNAs (Extended Data Fig. 6c). This result suggests that many unannotated RNAs, in particular regulatory elements derived RNAs, engage in genomic contacts that cannot be explained by a diffusion process around the transcription locus.

To better understand the nature of these contacts, we examined how far from the RNA transcription locus these contacts occurred (Fig. 5f). We found that most of the significant contacts made by UTL occurred within 100 kb of their locus (51% of all contacts), particularly for readthrough RNAs, which made over 69% of their contacts within 100 kb of their locus (Extended Data Fig. 6e). In contrast, introns of annotated RNAs showed deviations from the predicted patterns at larger distances. Indeed, only 17% of contacts from introns that were not predicted by the model occurred within 100 kb of their locus, whereas 88% occurred between 100 kb and 10 Mb. The difference in distances between RNA loci and their significant DNA contacts between annotated intron RNAs and unannotated RNAs suggests different types of interactions might be regulating RNA spread across chromosomes. Because these length scales are reminiscent of those involved in genome organization at the levels of TADs and A/B compartments^81–83^, we examined the relationship between the RNA localization patterns and the 3D organization of the genome.

### The 3D genome organization enables contacts between RNAs and distal chromatin loci

To examine how the 3D organization of the genome affects the localization patterns of individual RNAs on chromatin, we focused on a small ∼50 kb TAD on chr4q25, which is nested inside a larger 100 kb TAD (Fig. 6a). Two genes are located at the inner boundary of the small and large TADs: AC106864, an uncharacterized lncRNA, and the LARP7 gene, which is antisense to AC106864 and is highly transcribed in ES cells. We examined the binding profile of AC106864 on chr4 and found that most of the contacts of this RNA were within a few kb of its locus. We also observed two side peaks, labeled L1 and L2, that coincided with the other edge of the small and large TAD. In contrast, our generated model predicted a small peak at L1 (likely due to high accessibility of this locus as revealed by ATAC-seq) and no signal at L2. The fold difference signal of the observed data over the model confirmed that the two peaks at L1 and L2 were not explained by a simple diffusion of the AC106864 or accessibility biases. Interestingly, Hi-C data showed two corner peaks characteristic of a chromatin loop linking the LARP7 locus with both L1 and L2. This result suggests that AC106864 localization at L1 and L2 might be mediated by the chromatin loop. It is also possible that AC106864 targets these loci through other mechanisms such as base-pairing or association with RBP that are independent of genome folding. Yet this biochemically targeted interaction is unlikely given that the introns of the overlapping mRNA LARP7 also have contact peaks at L1 and L2. Together, these data suggest that TAD organization influences the contact patterns of RNAs, and that chromatin looping enables distal RNA-DNA interactions.

**Figure 6.**
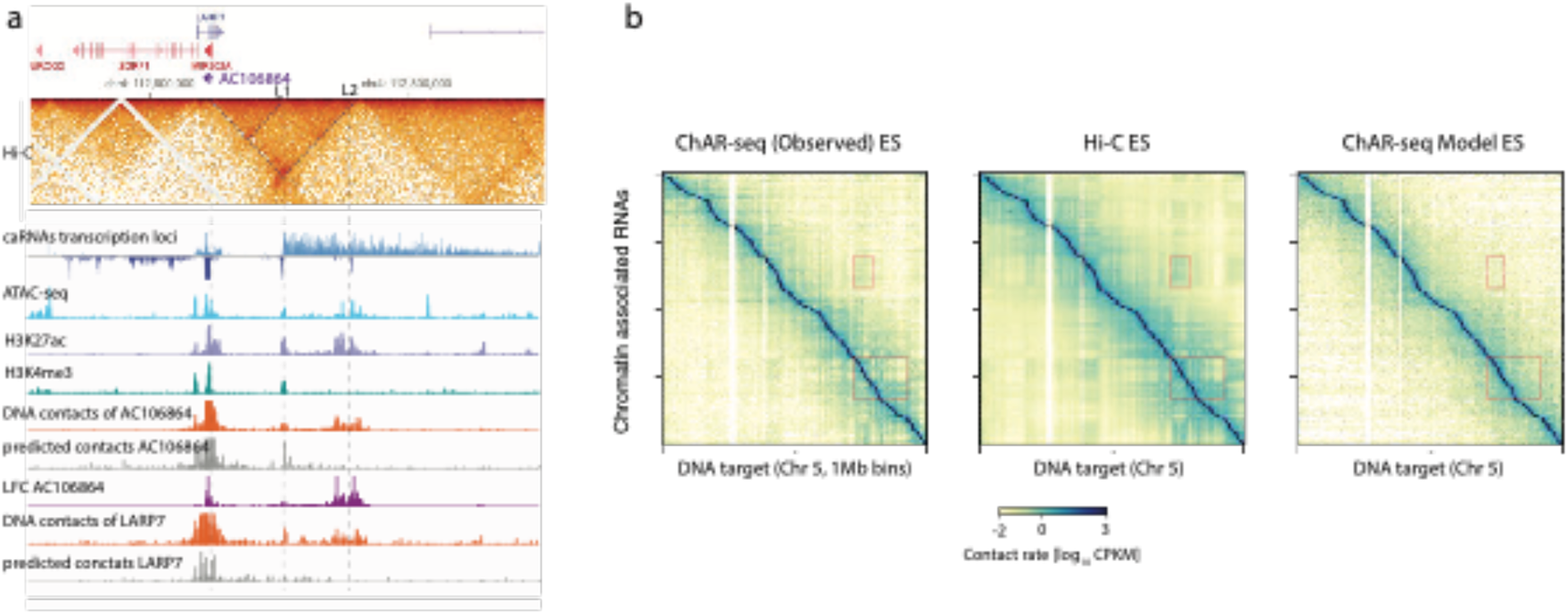
The 3D genome organization enables long-distance RNA-DNA contacts. a, Example of long-range RNA-DNA contacts across a chromatin loop at the LARP7 & AC106864 locus in ES calls. ICE normalized Hi-C map (2 kb resolution) is shown at the top. Transcription of LARP? (expressed from the positive strand) AC106864 (expressed from the negative strand, shown as negative values) are detected by ChAR-seq (top 2 tracks). The observed (dark organge) and predicted localization pattern (dark grey) of AC106864 on chromatin are shown with the log fate difference between observed and predicted (purple). The observed and predicted localization patterns for LARP7 are shown in light orange and light gray. AFAC-seq, H3K27ac and H3K4me3 tracks are also shown and indicate that L2 has enhancer-Ka chromatin properties. b, Comparison between ChAR-seq and Hi-C at the chromosome scale. Dashed Coves highlight two example regions where the WB. compartments Sad pattern io dearly visible in both Hi-C and ChAR-seq maps.

This observation prompted us to ask whether larger-scale topological organization of the chromosome also influences RNA-DNA contacts (Fig. 6b). ChAR-seq contact maps are naturally asymmetric in that the *y* -axis maps each row to an individual RNA and the *x* -axis maps each column to a genomic bin. To compare ChAR-seq to Hi-C data at the chromosome scale, we collapsed one dimension of the Hi-C maps into genes while keeping the other dimension as genomic bins. In these transformed Hi-C maps, each pixel represents the contact frequency between the gene and a cognate DNA bin. We detected in the ChAR-seq maps the same plaid pattern found in Hi-C data resulting from the 3D partitioning of the genome into two major compartments, the A and B compartments, also associated with active and inactive chromatin, respectively^83^. This pattern indicates that any individual caRNA tends to have a specific compartment (either A or B) with which it interacts preferentially. Equivalently, when one caRNA contacts a locus in say the A compartment, it has higher likelihood to contact other loci in the A compartment rather than in the B compartment. It was not surprising that this pattern was not produced by our generative model, since only linear distance is encoded in the model. We concluded that A/B compartments also modulate the long-range interactions of individual RNAs with chromatin.

### The caRNA-gene interactome preferentially links upregulated caRNAs to upregulated proximal genes

Our results point to a model where RNA-chromatin association patterns and their dynamics are restricted by i) the caRNA expression level ii) the genomic distance from the RNA locus to the DNA target, iii) the 3D chromatin topology. We wanted to determine whether this result is compatible with the hypothesis that ncRNAs participate in the regulation of cell-state specific protein-coding genes. We reasoned that RNAs with transcriptional regulatory roles are likely to be found near their cognate gene, where they could modulate local chromatin state, TF binding, RNA polymerase, or the activity of gene-proximal regulatory elements. This colocalization hypothesis is consistent with the better studied ncRNAs with gene regulatory activity, including XIST^17^, KCNQ1OT1^18^, and HOTAIR^84^. Thus, we defined a “proximal regulatory region” (PRR) around each protein-coding gene, encompassing +10 kb upstream and -90 kb downstream of its TSS, and measured the contact density of each caRNA at the PRR of each gene. Using this approach, we mapped all the physical contacts between the chromatin associated transcriptome and protein-coding genes (hereinafter referred to as the caRNA-gene interactome, Fig. 7a).

**Figure 7.**
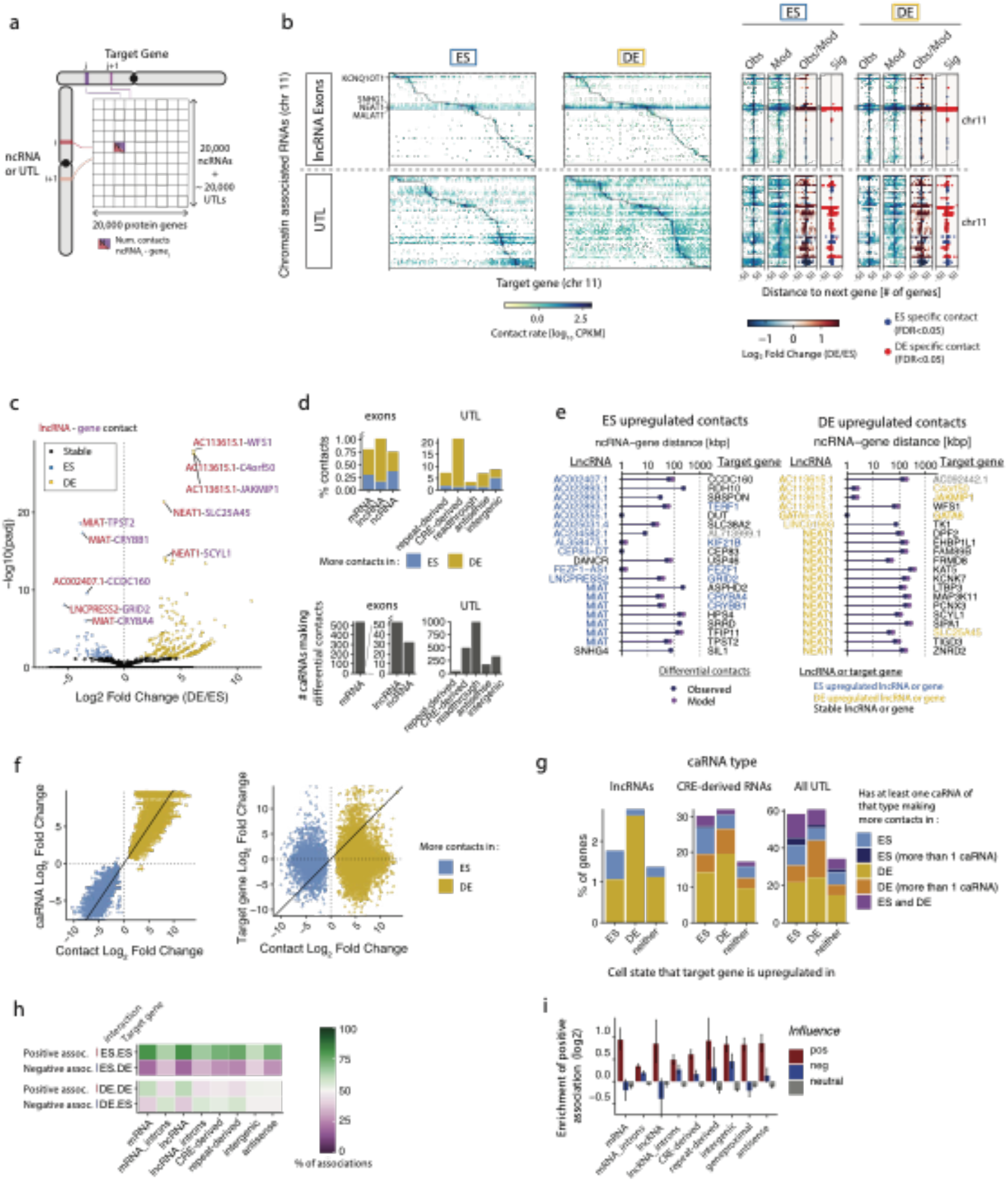
The caRNA-gene interactome preferentially links upregulated caRNAs to upregulated genes. **a**, Abstract representation of the caRNA-gene interactome studied in this analysis and displayed as a contact matrix with one caRNA per row and one protein-coding gene per column. Each matrix entry contains the number of contacts between an ncRNA and the proximal regulatory region (PRR) of a protein coding gene. Only *cis* interactions are shown for simplicity, but *trans* interactions are represented similarly. **b**, caRNA-gene interactome in ES and DE cells for the 50 most abundant lncRNAs (top) and UTLs (bottom) on Chr11. Narrow maps on the right are zoomed in view of the interactome for 50 protein-coding genes upstream and downstream of each caRNA PRR. Zoomed in maps are shown for the true interactome signal (Obs), the interactome predicted by the generative model (Mod), the log2 fold change (LFCobs over model) of the observed data over model (Obs/Model), and the interactions significantly enriched in the observed data over the model (Sig), namely those with adjusted *p*-value *<* 0.05 and LFCobs,model>0, obtained using DESeq2 as described in Fig. 4c-d. Only contacts with at least 10 counts in at least 2 samples were tested for enric hment over model. **c**, Volcano plot showing the differential lncRNA-gene contacts is ES versus DE cells. Each data point is a contact between a lncRNA and the PRR of a protein-coding gene. Volcano plots are shown as contact adjusted *p*-value versus log*2* Fold Change in DE versus ES cells (LFCES,DE). Differential contacts are computed as in Fig. 3a, and significant contacts are those with an adjusted *p*-value<0.05. **d**, Quantification of the percentage of cell-state specific contacts for each class of caRNA relative to the number of contacts tested for that class (top), and number of distinct caRNAs involved in these contacts (bottom). Cell-specific contacts were defined as those with an adjusted *p*-value<0.05 and LFCES,DE>1.3. **e**, Top 20 lncRNA-gene contacts upregulated in ES (left) and DE cells (right) in the observed data (blue circles). Most of these contacts are also predicted to be amongst the 20 most upregulated contacts by the generative model (purple circles). **f**, Scatter plots showing for each differential contact the relationship between the change in contact rate during differentiation (LFCES,DE) and the change in the chromatin levels of the involved caRNA (left) and in the expression of the cognate protein coding gene (right). Differential contacts were defined as in d). Only differential contacts involving exons of lncRNAs or UTLs are shown. **g**, Percent of protein-coding genes, targeted by one or more dynamic contact with a lncRNA (left panel), a CRE-derived RNA (middle panel), or any UTL (right panel, excluding tRNA- and snRNA-derived NARs). Protein coding genes are grouped (*x* -axis) according to whether their expression is upregulated in ES, DE, or stable during differentiation as measured by total RNA-seq (DEseq2, FDR 0.05, fold change threshold 3x). Colors indicate whether the protein coding gene is targeted by a single (light colors) or several (dark colors) caRNAs with which the interaction is upregulated in ES (blue shade) or DE (yellow shade). Some genes are targeted by several caRNAs which include both ES and DE upregulated interactions (purple). **h**, Top two rows: Percentage of interactions upregulated in ES targeting a protein coding gene upregulated in ES, which we define as a positive association, or targeting a protein coding gene upregulated in DE, which we define as a negative association. Bottom two rows: similarly, for interactions upregulated in DE cells. *x* -axis indicates the caRNA class. **i**, Fold enrichment of the fraction of positive associations in the observed interactome, compared to a randomized interactome, where the differential expression state of the target genes is shuffled. Error bars indicate 95% confidence intervals by bootstrap. Error bars not overlapping with *x* -axis indicate *p*-value <0.05 by bootstrap.

Consistent with the dynamics of the genome-wide RNA-DNA interactome (Fig. 3a-d), the caRNA-gene interactome of >1 million contacts was dynamic across differentiation. We detected most of the differential contacts at genes near the RNA locus (Fig. 7b). For lncRNAs only, we detected 340 differential contacts (∼1% of all lncRNA-gene contacts), but these involved only 57 distinct lncRNAs, indicating that a typical single lncRNA differentially contacts multiple genes (Fig. 7c,d). The caRNA-gene interactome involving UTLs was more dynamic than that involving annotated RNAs, consistent with the global interactome dynamics, with up to 20% differential UTL-gene contacts between ES and DE (Fig. 7d).

To identify potential regulatory caRNAs and their putative gene targets, we classified each caRNA and each protein-coding gene as an ES, DE, or stable caRNA or gene, based on those cells (FDR cutoff 0.05, Fold Change cutoff 3). We then examined the statistical associations between the class (ES/DE/stable) of a caRNA, its cognate gene, and their interaction. Fig 6e shows the top 20 most upregulated contacts involving a lncRNA along with the cognate lncRNA-gene pair. We noted that all the top 20 upregulated contacts in a given cell state involved ncRNAs upregulated in the same state. This result is consistent with our findings that the RNA-DNA interactome dynamics is globally driven by transcriptional dynamics. Yet most of the nearby genes for these differential contacts were not differentially expressed in ES vs DE, suggesting that changes in the caRNA levels at these genes do not affect their expression. Furthermore, the fold change in contact rate during ES to DE transition correlated with the fold change of the expression of the source caRNA (Fig. 7f, left panel), but not with that of the contacting protein coding gene (Fig. 7f, right panel).

To further understand the relationship between gene expression and presence of a caRNA in the PRR of a gene, we examined how many cell-state specific contacts are made at cell state-specific genes. This analysis revealed that >97% of cell state specific genes are not contacted by lncRNAs in a cell state specific manner (Fig. 7g, left panel). Interestingly however, over 50% of these genes are contacted by at least one, and sometimes several UTL specifically in one cell state (and 15% with a CRE). In contrast, only ∼25% of genes that are not cell-state specific were contacted by cell-state specific UTLs. Thus, most genes do not require cell-state specific localization of a particular lncRNA in their PRR to alter their expression, but genes whose expression is altered are likely to be contacted by an UTL in a cell-state specific manner.

Together, our findings indicate that the presence of an individual ncRNA near the gene TSS does not correlate with the gene’s transcription. This result does not rule out a regulatory activity of ncRNAs at protein coding genes. It remains possible that multiple inputs gate the target gene’s expression, including chromatin state, transcription factors, and possibly several RNAs, which could wash out average correlations between caRNA-gene interactions and gene transcription.

To identify patterns in the interactome that could reveal a regulatory structure, we compared the observed interactome dynamics to that which would be expected should it be independent of the gene expression dynamics (null model). We binned differential contacts in 3 categories: i) positive edges, where the contact dynamics were positively correlated with the proximal gene dynamics (contacts that increased in ES to genes that increased in ES, or contacts that increased in DE to genes that increased in DE), ii) negative edges (contacts that increased in ES to genes that increased in DE, or contacts that increased in DE to genes that increased in ES), iii) neutral edges (contacts that increased in ES or DE to genes that were neither ES or DE genes).

We found that across all categories of caRNAs, the interactome contained up to 1.8 times more positive edges (*p*-value<0.05 by bootstrap) and up to 1.3 times fewer negative edges (*p*- value<0.05 by bootstrap) than would be expected for a random interactome under the null model (Fig. 7h,i). Thus, we conclude that although specific RNAs are not sole drivers of transcription activation or silencing at any gene, the architecture of the interactome is consistent with an overall positive regulation, where the presence of caRNAs is generally associated with higher expression of the contacted genes.

## Discussion

Understanding how caRNAs control chromatin state and transcription is a long-standing problem. To date, only a few RNAs have been linked to specific regulatory functions. In this work, we provide a global view of the RNA-chromatin interactome that expands studies focused on individual RNAs and uncovers general principles governing the architecture of putative ncRNA-gene regulatory networks.

First, we show that lncRNAs with promiscuous chromatin interactions are rare. Given that we detected only a handful of lncRNAs with such patterns, it is unlikely that uncharacterized lncRNAs have global regulatory roles, such as those established for 7SK, MALAT1, XIST, or TERC. However, we identified a larger repertoire of unannotated RNAs with broad chromatin interactions which contained many TE-derived RNAs. These data reinforce the idea that transcriptionally active LINE, SINE and LTR may play key roles in chromatin regulation and highlight the necessity to further explore the biology of transposable elements^85 84^.

Second, it is noteworthy that all delocalized lncRNAs but TERC and 2 uncharacterized ncRNAs (VAULT-RNA and AC073335) are known RNA residents of the nucleolus (RMRP, RPPH1, 7SL, most snoRNAs) or nuclear speckles (7SK, MALAT1, most snRNAs). SPRITE, a Hi-C-like method which probes high order chromatin interactions, showed that the 3D genome is organized into 2 major hubs around the nucleolus and nuclear speckles, where abundant long-range and interchromosomal DNA-DNA contacts occur^86^. We hypothesize that the proximity of these RNA loci to the genomic hubs may be important in enabling interactions with dispersed genomic loci. This behavior is reminiscent of XIST, whose location on the X chromosome defines where heterochromatin spreading initiates^17^. We speculate that a general principle may underlie these observations, where the interactions of an RNA with chromatin are constrained by the position of their transcription locus relative to other loci or to nuclear domains.

Third, we demonstrate unambiguously that no RNA behaves like XIST and localizes throughout its own chromosome while being excluded from other chromosomes. Thus, while XIST sets expectations for ncRNAs regarding their potential roles as regulators of transcription and large genetic networks, XIST appears to be unique in its localization pattern.

Fourth, excluding small RNAs such as snRNAs, snoRNAs, tRNAs, and some UTLs as described above, we found no evidence across the non-coding transcriptome for widespread existence of *trans* interactions, or of affinity-driven interactions as previously defined^2^. Indeed, we demonstrated that a simple generative model, encoding only for expression and RNA-DNA distance, accurately predicts the contact patterns for each RNA. Thus, while acknowledging possible false-negatives for lowly expressed RNAs, we show that nearly all interactions are proximity-driven. Our data does not differentiate between RNAs that are tethered to chromatin via active transcriptional complexes versus other mechanisms such as nucleoprotein complex interaction or base pairing. We anticipate that multiple different mechanisms may act to retain RNAs in proximity to their sites of synthesis. An important implication of our results is that across the non-coding transcriptome, chromatin regulatory activities are essentially limited to nearby genes. Our observations are consistent with previous studies of lncRNAs that demonstrate a propensity for localization near the transcriptional locus and cis-regulation of gene activity ^58, 87–89^.

Several modes of regulatory activity are compatible with proximity-driven interactions, yet our work brings in important refinements to the proposed models. If a ncRNA serves as a platform to locally recruit histone modifying complexes, as proposed for many lncRNAs, we show that the dimensions of the domain around the RNA transcription locus where this activity occurs is solely determined by the RNA expression. The same local constraints apply if a ncRNA operates via a decoy mechanism, whereby it evicts specific remodeling complexes from chromatin, through competitive or inhibitory associations with these complexes^90^.

To our surprise, we observed a general lack of correlation between the dynamics of the RNA contacts at a given gene and the dynamics of the expression of that gene. This observation challenges models proposing that the activation or silencing of a gene may be controlled by a single ncRNA^18, 23, 77, 84, 91^. Instead, our data indicates that most ncRNAs do not have gene regulatory activity, or favors some of the more complex proposed models, for example, involving coordinated inputs from a ncRNA and the local chromatin environment. One such model, the “junk mail model,” posits that caRNAs interact with chromatin remodeling complexes and keep them poised and in check until other local conditions are satisfied^48, 92^ (such as deposition of a specific chromatin modification or binding of a transcription factor). The junk mail model is compatible with our observations. Another possibility, which we termed the “democratic RNA model,” is that the distributed activity of multiple, weakly influential ncRNAs, rather than that of a single, strongly influential ncRNA determines the overall regulatory output of RNA-chromatin interactions at a gene. The lack of a strong correlation between RNA interaction and gene expression indicates that testing the functions of individual RNAs in gene regulation may be challenging and evaluating RNA regulatory roles will require combinatorial perturbations of RNAs and putative effectors at specific loci.

We found that an increase in interaction frequency between a specific ncRNA and a target gene is more likely to correlate with an increase in target gene expression than one would expect should the ncRNA-gene contacts and the gene expression be uncorrelated with one another.

Three scenarios may explain this result. First, this may merely reflect increased accessibility during chromatin activation and higher likelihood to crosslink nearby RNAs. Second, there may be local coregulation of nearby ncRNAs and genes, for instance, through shared regulatory elements. Third, it is possible that the default activities of caRNAs are: i) a decoying of the silencing machinery, as proposed by the junk mail model, in the context of PRC2 eviction^93^, or ii) a recruitment of transcription activators such as the CREB-binding protein^30^. These two effects would also give rise to a positive correlation between caRNA presence at a gene and transcriptional output of this gene.

As mentioned previously, we did not identify lncRNAs localized at defined genomic targets in *trans*, beyond the interactions explained by the expression levels of these lncRNAs and the distance to their targets. This finding will need to be reconciled with the models proposed for a few ncRNAs, such as DIGIT or RMST, which have been reported to broadly colocalize with BRD3 at endoderm differentiation genes, and SOX2 at genes that control pluripotency and neurogenesis, respectively^77, 78, 91^. Given that lncRNAs and eRNAs are highly cell state specific^33, 34^, the architecture of the caRNA-chromatin interactome may be qualitatively different and perhaps contain more *trans* interactions in further differentiated cells. Additionally, we cannot exclude the possibility that our analysis missed affinity-driven trans interactions due to sequencing depth limitations, in particular for lowly expressed ncRNAs. Thus, deeper sequencing or more powerful statistical frameworks may reveal weak deviations from the model at more loci. However, the fact that our analysis reveals broad differences in the contactome between ES and DE cells gives us confidence that any undetected deviation from the model must be more subtle than the contactome changes related to cellular differentiation.

This work presents a global analysis of caRNA-chromatin interaction and establishes that caRNAs predominately operate locally, through diffusion and genome conformation driven interactions. We anticipate this work will direct the efforts in the non-coding RNA field by providing data-informed priors on the localization of RNAs, and a simple model predicting where non-coding RNAs may act. Future studies to identify the proteins mediating these RNA-chromatin interactions will be necessary to inform the interplay between caRNA and RNA binding proteins in the control of transcription and chromatin state.

## Methods

### Human H9 ES cell culture

H9 hESCs cells (ES cells) were obtained from Wicell (cell line WA09) and cultured on Corning Matrigel hESC qualified matrix with mTeSR1 medium according to manufacturer’s protocols and as described in Loh et al. 2014^45^. Briefly, 6-well plates were prepared with matrigel by adding 1 mL of matrigel (diluted in serum free DMEM/F-12 according to lot dilution factor) to each well and polymerized for 1 hour at room temperature. DMEM/F-12 was aspirated and replaced with 1.5 mL mTeSR1 warmed to room temperature, then 2 *µ*M of 10 mM ROCK inhibitor (Y27632- Dihydrochrolride) was added to each well. H9 hESCs (∼3-5 million cell aliquots) were thawed and immediately diluted by dropwise addition of 10 mL prewarmed mTeSR1, spun at 200 g for 5 min, and gently resuspended in 1.5 mL mTeSR. 0.5 mL of cells were added to each well and placed at 37 °C. Media was replaced daily with 2 mL fresh mTeSR1 per well. When colonies were ∼70% confluent and started to touch each other, cells were passaged as colonies. Each well was washed with 1x PBS, 1 mL of Versene-EDTA was added, and cells were incubated at 37 °C for 5 min. Colonies were detached, broken up with gentle pipetting, and resuspended in mTeSR1 at a 1:5 to 1:10 dilution. 0.5 mL of cells was added dropwise to each well containing 1.5 mL mTeSR1 and coated with matrigel prepared as described above (without ROCK inhibitor).

### Differentiation into definitive endoderm

Colonies were seeded from 1:10 dilution on day 0 into four 15 cm dishes with matrigel, two for maintenance as ES cells and two for differentiation into Definitive Endoderm (DE) cells. ES cells were maintained as above with daily mTeSR1 media replacement. For differentiation, cells were treated with 10 *µ*M ROCK inhibitor on day 0, and their media was replaced on day 1 with DE induction Media A (Gibco Cat# A3062601), and on day 2 with DE induction Media B (Gibco Cat# A3062601). On day 3, cells were harvested for ChAR-seq. In addition to the 15 cm dishes used for ChAR-seq, cells were also seeded and maintained as ES cells or differentiated into DE cells in 6-well plates with poly-L-lysine coated coverslips under matrigel, and collected at the same time for immunofluorescence analysis. Cells were also differentiated in 6-well plates for RNA-seq and ATAC-seq.

### Immunostaining

Cells were cultured in 6-well plates on poly-L-lysine coated coverslips under matrigel and maintained as ES cells or differentiated into DE cells as described above. Cells were washed three times with PBS and fixed with 2% PFA in PBS added directly to the wells for 10 min at room temperature. The PFA solution was aspirated, cells were washed three times with PBS, and permeabilized with 0.1% Triton-X-100 in PBS for 5 min at room temperature. Coverslips were transferred to parafilm-coated staining chambers, washed with PBS, and blocked with Antibody Dilution Buffer (AbDil, 150 mM NaCl, 20 mM Tris-HCl pH 7.4, 0.1% Triton X-100, 2% BSA, 0.1% Sodium Azide) for 30 min at room temperature. Samples were incubated in primary antibody for 30 min at room temperature (Rabbit anti-Nanog 1:500, Goat anti-Sox17 1:1000 diluted in AbDil), washed three times with AbDil, and incubated with secondary antibodies conjugated to Goat anti-Rabbit Alexa-647 and Chicken anti-Goat Alexa-555 (1:1000 diluted in AbDil) for 30 min at room temperature. Cells were washed with AbDil three times, stained for 5 min with 10 *µ*g/mL Hoechst-33342 in PBS, and washed with PBS with 0.1% Triton-X-100 before being mounted (20 mM Tris-HCl pH 8.8, 0.5% p-Phenylenediamine, 90% glycerol) onto slides and sealed with nail polish. Samples were imaged with an IX70 Olympus microscope with a Sedat quad-pass filter set (Semrock) and monochromatic solid-state illuminators. Cells were imaged using a 40x objective. At least 10 images per coverslip were captured using 0.2-*µ*M *z*- stacks. Maximum intensity projections were processed with CellProfiler (3.1.8) to identify nuclei based on Hoechst signal and to measure the mean intensity of each channel. Histograms of mean nuclear intensity for each marker were plotted in R.

### qPCR

For qPCR, RNA was extracted from each well of a 6-well plate containing ES or DE cells (∼1 million cells per well) using 1 mL Tripure reagent and according to the manufacturer’s protocol. RNA were treated with DNase (TURBO DNase; Ambion) for 1 hour at room temperature followed by isolation with a minElute RNA Cleanup Kit (Qiagen). RNA concentrations were measured by Nanodrop and total RNA integrity assayed using an Agilent Bioanalyzer. All RNAs had a RNA integrity number (RIN) greater than 9.0. 0.5-1 *µ*g of RNA was reverse-transcribed with random hexamer primers using SuperScript III reverse transcriptase (18080-051; Invitrogen) according to the manufacturer’s protocols. First-strand cDNA was diluted 1:10 in nuclease free H2O and amplified using gene-specific primers that had been tested for amplification efficiencies >90% and to amplify a single product. Real-time PCR was performed using the Powerup SYBR Master Mix (ThermoFisher) for 40 cycles (94 °C 15 sec, 55 °C 30 sec, 68 °C, 1 min) on an ABI ViiA 7 Real-Time PCR Machine with cycle thresholds (CTs) determined automatically and with all samples in triplicate. Experimental genes were normalized to the PBGD housekeeping gene, with relative expression levels calculated using the 2^ΔΔ^*^CT^* method, and the transcript level fold-change in DE versus ES cells was calculated. If a gene’s expression was too low to detect via qPCR, these “undetermined” Ct values were assigned a value of 38 to provide a conservative over-estimate for use in calculation of expression change.

### RNA-seq

For RNA-seq, RNA was extracted from each well of a 6-well plate containing ES or DE cells using 1 mL Tripure and the Direct-Zol RNA Extraction kit (Zymo Research) according to the manufacturer’s instructions. RNA concentrations and quality were assayed as described for qPCR. For each sample, 2.5 *µ*g of RNA was treated with DNase (TURBO DNase; Ambion) for 1 hour at room temperature followed by isolation with a RNA Clean & Concentrator-25 kit (Zymo Research). 1 *µ*g RNA was converted to ribosomal depleted cDNA libraries ready for sequencing using the TruSeq Stranded Total RNA Library Prep Human/Mouse/Rat kit (Illumina) according to the manufacturer’s instructions. Samples were uniquely dual indexed using IDT for Illumina TruSeq RNA UD Indices. The 4 biological replicates from both conditions (ES and DE cells) were pooled and sequenced at low read depth on a MiSeq (2 x PE75) at the Stanford Functional Genomics Facility to assess quality, and later on 1 lane of the HiSeq4000 (2 x PE150) at NovoGene (Sacramento, CA). All reported analysis was generated using the HiSeq dataset.

### ATAC-seq

Cells for ATAC-seq were differentiated as described above and collected by dissociating in Versene followed by resuspension in warm mTeSR media. Cells were transferred to 15 mL conical tubes and centrifuged at 1000 RPM for 5 min. The pellet was resuspended in DPBS, cells were counted and immediately processed. ATAC-seq was performed as previously described using the OMNI-ATAC protocol (Corces et al., Nature Methods 2017) with slight modifications. Briefly, ∼100K cells were resuspended in 50 *µ*L cold ATAC-Resuspension Buffer (10 mM Tris-HCl pH 7.4, 10 mM NaCl, 3 mM MgCl2, 0.01% Digitonin, 0.1% Tween-20, and 0.1% NP40 in water) and incubated on ice. Cells were washed with 1 mL cold ATAC-RSB (without NP40 and digitonin), centrifuged at 500 RCF for 10 min at 4 °C. The pellet was resuspended in 50 *µ*L transposition mixture (2x TD buffer and 2.5 *µ*L transposase) from the Illumina Nextera DNA Library Prep Kit and incubated at 37 °C for 30 min in a thermomixer at 1000 RPM. Libraries were purified with the DNA Clean & Concentrator-5 Kit (Zymo Research) and PCR amplified with barcoded primers (Buenrostro et al., 2013). Amplification cycle number for each sample was monitored by qPCR to minimize PCR bias. PCR amplified libraries were purified with the MinElute purification kit (Qiagen) and excess primers and large (>1000 bp).

DNA fragments were removed by AMPure XP bead selection (Beckman Coulter). Four biological replicates from each cell type (ES and DE) were pooled and sequenced at low read depth on a MiSeq (2 x PE75) at the Stanford Functional Genomics Facility to assess quality and later on 1 lane of the HiSeq4000 (2 x PE150) at NovoGene (Sacramento, CA). All reported analysis was generated using the HiSeq dataset.

### ChAR-seq library preparation

ChAR-seq libraries were prepared according to the published protocol^42^ as briefly described below. All reagents used were RNAse-free.

### Cell fixation and nuclei

About 10 million cells were harvested from a 15 cm dish with Versene and fixed in 3% formaldehyde for 10 min at room temperature. Formaldehyde was quenched with the addition of 0.6 M glycine for 5 min at room temperature then 15 min on ice. Cells were pelleted for 5 min at 500 g at 4 °C, washed with 10 mL ice-cold PBS, and resuspended in ∼5-10 mL PBS. Cell concentration was measured and cells were aliquoted in batches of 10 million cells in 1.5 mL tubes. Aliquots were spun for 5 min at 500 g at 4 °C, the supernatant was removed, and pellets were flash frozen in liquid nitrogen and stored at −80 °C until library preparation.

### Cell lysis and nuclei preparation

Frozen pellets were resuspended in 500 *µ*L ice cold lysis buffer (10 mM Tris-HCl pH 8, 10 mM NaCl, 0.2% Igepal-CA 630, 1 mM DTT, 1 U/*µ*L RNaseOUT, 1x protease inhibitor) and incubated for 15 min on ice. Nuclei were washed (throughout the protocol, “nuclei were washed” indicates the following steps: spinning for 4 min at 2500 g, discarding of supernatant, resuspension and mixing in the indicated wash buffer, spinning for 4 min at 2500 g, and aspiration of the wash buffer) with 500 *µ*L of lysis buffer without Igepal, RNaseOUT, or Protease Inhibitor, then resuspended in 400 *µ*L of 0.5% SDS (10 mM Tris-HCl pH 8, 10 mM NaCl, 1 mM DTT, 0.5% SDS, 1 U/*µ*L RNAseOUT), and incubated for 10 min at 37 °C. SDS was then quenched by adding Triton X-100 to 1.4% final concentration and incubating for 15 min at 37 °C.

### In situ biochemistry steps for RNA-DNA proximity ligation

To fragment RNAs, nuclei were pelleted and resuspended in 150 *µ*L fragmentation buffer (0.25x T4 RNA ligase buffer, 1 U/*µ*L RNAseOUT), and exposed to heat for 4 min at 70 °C. To dephosphorylate RNA 5’ ends, nuclei were washed twice (in 800 *µ*L PBS then 800 *µ*L 1x RNA ligase buffer, with the first spin omitted for the first wash and PBS added directly to the previous reaction), resuspended in 150 *µ*L dephosphorylation mix (1x T4 PNK buffer, 1 U/*µ*L T4 PNK, 1 U/*µ*L RNAseOUT), and incubated for 30 min at 37 °C. To perform RNA-bridge ligation, nuclei were washed twice as above and resuspended in 200uL RNA-bridge ligation mixture [1x T4 RNA ligase buffer, 25 *µ*M annealed ChAR-seq bridge (top strand: /5rApp/AANNNAAACCGGCGTCCAAGGATCTTTAATTAAGTCGCAG/3SpC3/; bottom strand: /5Phos/GATCTGCGACTTAATTAAAGATCCTTGGACGCCGG/iBiodT/T; individual strands ordered from IDT DNA), 10 U/*µ*L T4KQRNAligase2, 1.5 U/*µ*L RNAseOUT, 20% PEG-8000] and incubated overnight at 23 °C on a thermomixer at 900 RPM. To perform first-strand synthesis, nuclei were washed twice as above and resuspended in 250 *µ*L of first strand synthesis mixture (1x T4 RNA ligase, 8 U/*µ*L Bst3.0, 1 mM of each dNTP, 1 mM DTT, 1 U/*µ*L RNAseOUT), and incubated for 15 min at 23 °C, 10 min at 37 °C, and 20 min at 50 °C. Bst3.0 was inactivated by adding 8 *µ*L of 0.5 mM EDTA (15 mM final concentration), 14 *µ*L of 1% SDS (0.5% final), and incubating for 10 min at 37 °C. SDS was then quenched with 43 *µ*L of 10% Triton X-100 (1.3% final concentration) for 15 min at 37 °C. Next to perform genomic digestion, nuclei were washed twice and resuspended in 250 *µ*L of DpnII reaction mixture (1x T4 RNA ligase, 3 U/*µ*L DpnII, 1 mM DTT, 1 U/*µ*L RNAseOUT) overnight at 37 °C on a thermomixer at 900 RPM. DpnII was inactivated in the same manner as Bst3.0 inactivation. SDS was quenched as above. Next, to perform bridge-DNA ligation, nuclei were washed twice and resuspended in 250 *µ*L of ligation mixture (1x T4 DNA ligase, 10 U/*µ*L T4 DNA ligase, 1 U/*µ*L RNAseOUT) for 4 hours at 23 °C. T4 was inactivated by adding 8 *µ*L of 0.5 M EDTA (15 mM final concentration). Finally, to perform second strand synthesis, nuclei were washed twice (PBS then 1x cDNA buffer 10 mM Tris-HCl pH 8, 90 mM KCl, 50 mM (NH4)2SO4), and resuspended in 250 *µ*L of secon strand synthesis mix (1x cDNA buffer, 0.5 U/*µ*L *E. coli* DNA PolI, 0.025 U/*µ*L RNaseH, 1 mM of each dNTP, 1 mM DTT) for 1.5 hours at 37 °C.

### DNA isolation and shearing

Reverse crosslinking was carried out by adding 31.25 *µ*L of 10% SDS, 31.25 *µ*L 0.5 M NaCl, 9 *µ*L of 20 mg/mL proteinase K and incubating overnight at 68 °C. DNA was purified by phenol chloroform extraction, ethanol precipitated, and resuspended in 130 *µ*L TE (10 mM Tris pH 8, 0.1 mM EDTA) buffer. DNA was sheared with a Covaris S220 to target a mean fragment size of ∼200 bp (175 peak incident power, 10% duty factor, 200 cycles/burst, 180 sec). Fragment size distribution was quality controlled on an Agilent High Sensitivity DNA Bioanalyzer.

### Isolation of biotinylated molecules, on-beads adapter ligation, and on-beads PCR

Molecules containing the biotinylated bridge sequence were isolated using 150 *µ*L of MyOne Streptavidin T1 dynabeads. To bind bridge containing molecules, beads were washed with 750 *µ*L tween wash buffer (TWB, 10 mM Tris pH 8, 0.5 mM EDTA, 1 M NaCl, 0.05% Tween20) and resuspended in 130 *µ*L 2x bead binding buffer (10 mM Tris pH 8, 2 M NaCl, 0.5 mM EDTA) and 130 *µ*L sheared DNA sample, then incubated at room temperature for 15 min with agitation. To remove unbound DNA, beads were washed twice with 750 *µ*L TWB (with incubation at 50 °C for 2 min with agitation during the first wash), then resuspended in 40 *µ*L TE buffer. DNA ends were prepared for ligation by adding 7 *µ*L of NEBNext End Prep Buffer and 3 *µ*L NEXext End Prep enzyme mix and incubating for 20 min at room temperature and 30 min at 65 °C. Adapters were ligated using NEBNext Ultra II Ligation module according to manufacturer’s protocols. Beads were washed twice as above and resuspended in 50 *µ*L PCR amplification mix (25 *µ*L 2x NEBNext High Fidelity master mix, 2.5 *µ*L 10 *µ*M Universal Primer, 2.5 *µ*L 10 *µ*M indexing primer, 20 *µ*L H2O). PCR reaction was performed using the following program (1 cycle: 98 °C for 30 sec; 5 cycles: 98 °C for 10 sec, 65 °C for 75 sec). Beads were magnetically collected and the supernatant containing amplified DNA was transferred to a clean 1.5 mL microcentrifuge tube.

The amplified libraries were purified using magnetic SPRI beads at a ratio of 1:1 and eluted with 31 *µ*L 10 mM Tris-HCl, pH 8.

### Side qPCR & off-bead PCR

To determine the number of additional cycles of PCR amplification to perform, 5 *µ*L of purified library from on-bead PCR, 6 *µ*L 2x NEBNext High Fidelity master mix, 0.5 *µ*L 10 *µ*M Universal primer, 0.5 *µ*L 10 *µ*M indexing primer, and 0.33 *µ*L 33x SYBR Green were mixed and added to a qPCR well and cycled on an ABI ViiA 7 Real-Time PCR Machine with the following parameters (1 cycle: 98 °C for 30 sec; 25 cycles: 98 °C for 10 sec, 65 °C for 75 sec). The number of off-bead PCR cycles to perform was determined by finding the number of cycles such that the fluorescence intensity is about one third the plateau intensity at the PCR saturation. The remaining 25 *µ*L of the library was combined with 30 *µ*L 2x NEBNext High Fidelity master mix, 2.5 *µ*L 10 *µ*M universal Primer, and 2.5 *µ*L 10 *µ*M indexing primer. Each sample was then cycled as above for the number of cycles determined by the side qPCR.

### Library clean up and sequencing

To purify the amplified library, high molecular weight fragments were bound to Ampure beads by adding 0.6x volume of the PCR reaction of Ampure beads and collecting the supernatant. Low molecular weight fragments were purified out by adding 0.1875x the volume of the supernatant transferred to obtain a final ratio of 0.9x beads:slurry. DNA was eluded in 33 *µ*L 10 mM Tris pH 8. Library concentration was assessed using a Qubit dsDNA High Sensitivity kit and size distributions were determined using an Agilent High Sensitivity DNA bioanalyzer. Samples were pooled and sequenced on 1 lane of an Illumina HiSeq4000 platform (2x PE150) to assess library quality, then later deeply sequenced on 2 lanes of an Illumina NovaSeq platform at NovoGene (Sacramento, CA). All reported analysis was generated using the NovaSeq dataset. Replicates 1 and 2 of ES and DE ChAR-seq libraries were prepared at different times and each sequenced separately on 1 NovaSeq lane.

### ChAR-seq data processing and generation of pairs files

Demultiplexed fastq files from the ChAR-seq data were processed using a custom Snakemake pipeline (https://github.com/straightlab/charseq-pipelines), outputting pairs files containing the RNA and DNA coordinates of each RNA(cDNA)-DNA chimeric read and relevant annotations for each RNA-DNA contact. A summary of the pipeline workflow is depicted in Supplementary Fig. 1. For full details of the processing pipeline, see Supplementary Note 1. Briefly, reads were PCR deduplicated using clumpify.sh v38.84 (BBMap suite), low quality reads (*Q <* 30) were removed, and sequencing adapters were trimmed using Trimmomatic v0.38. Paired-end reads were merged using Pear v0.9.6 when possible and reads containing a single instance of the ChAR-seq bridge sequence were identified using chartools v0.1, a custom ChAR-seq reads preprocessing package released as part of this study (https://github.com/ `straightlab/chartools`). Reads were split into a rna.fastq and dna.fastq file corresponding to the sequences of the RNA (cDNA) and DNA side of the chimeric molecule using chartools. Reads with either the RNA or DNA side shorter than 15 bp were removed using chartools, and reads whose RNA side aligned to a rRNA sequence by Bowtie2 were filtered out using Picard. DNA reads were aligned to hg38 using Bowtie2, and RNA reads were aligned to hg38 using STAR and Gencode v29 annotations. RNA reads were assigned specific genes using tagtools (https://github.com/straightlab/tagtools), a package released as part of this study. Genes were assigned an RNA type (either mRNA, lncRNA or ncRNA) based on the GencodeV29 “gene type” field and the lookup table in Table S7, which we used to simplify the original Gencode classification. ncRNA genes were assigned a subtype, as indicated in Table S7, to break down this group into functional classes. Pairs files containing for each read the mapping coordinates of the DNA, the RNA, and the most likely gene of origin were produced using chartools `pairup` function. Separate pairs files were produced for reads whose RNA was annotated by tagtools as exonic, intronic, or intergenic. pairs files were filtered using a bash script to remove multimapping reads and reads with low mapping scores on either the RNA (STAR *Q <* 255) or DNA (Bowtie2 *Q <* 40) side. Reads whose RNA overlapped with the hg38 ENCODE blacklist or that could not be attributed to a single known gene or genomic locus were also removed.

### RNA-seq data processing

RNA-seq reads were processed using a Snakemake pipeline mirroring the ChAR-seq pipeline, but all of the operations related to the DNA-side of the reads were skipped. In brief, demultiplexed fastq files were deduplicated, sequencing adapters were removed, paired mates were merged as described for ChAR-seq reads. Reads that aligned to a rRNA sequence by Bowtie2 were filtered out using Picard. Reads were aligned to hg38 using STAR and were annotated with tagtools using the Gencode V29 gene models. Reads with low mapping scores (STAR *Q <* 255), reads which could not be attributed to a single known gene or a single locus, and reads that overlapped with a locus on the ENCODE black were discarded.

### ATAC-seq data processing

Illumina Nextera Adapters were removed using a custom Python script. Reads were aligned to the hg38 using Bowtie2. Duplicates were removed with Picard. Mitochondrial reads or reads with Bowtie2 MAPQ score *<* 30 were removed using SAMtools. All replicates were similar, so their alignment files were merged to increase library complexity (>100 million mapped reads per cell type) and produce a single bigwig file per cell type used to display the ATAC-seq tracks, and a single bam file to determine ATAC-seq peaks. ATAC-seq peaks were identified in each cell line using HMMRATAC v1.2.10.

### Chromatin association scores

We defined the chromatin association score for RNA *i* as the log fold difference between the level of RNA *i* in the chromatin associated RNA transcriptome (measured with the RNA-side of the ChAR-seq reads) and its level in the total RNA transcriptome (measured with total RNA-seq). To estimate the chromatin association score in a way that was robust to small counts and obtain *p*-values to detect RNAs with meaningful chromatin enrichment, we used DEseq2 with a design formula `∼cell + sequencing + cell:sequencing`. In this design matrix, the `cell` covariate represented the cell type and the `sequencing` covariate indicated whether the sample originated from RNA-seq or ChAR-seq. The interaction term `cell:sequencing` captured differences in the chromatin association of a given RNA between ES and DE cells. We used the shrunken estimate of the regression coefficient associated with the `sequencing` covariate as the estimate of the chromatin association score. We computed the chromatin association score in ES and DE cells separately by setting the reference level for the `cell` covariate to ES and DE, respectively, before running DEseq2. The apeglm method was used to compute the shrunken estimates. We ran DEseq2 using an input count matrix with 16 samples: 2 ES and 2 DE replicates from ChAR-seq and 4 ES and 4 DE replicates from RNA-seq. Gene counts for all Gencode V29 genes and all UTLs identified in this study were included in the input matrix, except those with fewer than 10 counts combined across all 16 samples.

Counts from exons and introns of a given gene and from UTLs were input as separate entries (rows) in the matrix. All DESeq2 parameters were set to their default value, except for the sample depth normalization step. For sample depth normalization, we ran the `estimateSizeFactors` command on a subset of the rows of the count matrix that included only exons of annotated genes with at least 50 counts combined across all 16 samples.

Subselecting exonic reads removed length bias due the low representation of introns in the total RNA-seq data compared to the ChAR-seq data. False Discovery Rate (FDR) adjusted *p*-values corresponding to the regression coefficient associated with the `sequencing` covariate were used to identify genes with significant chromatin enrichment. Genes with an adjusted *p*-value smaller than 0.05 and a chromatin association score either greater than 3 where labeled as chromatin enriched, and those with an adjusted *p*-value smaller than 0.05 and a chromatin association score less than -3 were labeled as chromatin depleted. To identify genes with statistically significant changes in their chromatin association score in ES versus DE cells (Fig. 3d), we used the regression coefficient associated with the interaction term `cell:sequencing`, LFCES,DE and its corresponding adjusted *p*-value *p*adj,ES,DE. Thresholds used to label such genes where LFCES,DE *>* 0, and *p*adj,ES,DE *<* 0.05.

### Computational interaction with ChAR-seq data

For most computational analyses, the filtered pairs files were loaded in python as a chartable python object using the chartools package. Within the object, the interaction data were stored in a sparse matrix with one row per RNA and one column per genomic DpnII site, binned at 10 bp resolution, which could be loaded entirely in RAM. This allowed us to perform computationally efficient indexing operations to select individual RNAs or target genomic loci, plot ChAR-seq maps at various resolutions, produce bigwig files of the binding profile of individual RNAs, and generate the caRNA-gene interactome. All of these operations were performed using methods from the chartools package. De-novo identification and classification of unannotated transcribed loci (UTLs)

### Identification of UTLs

For each ChAR-seq sample, reads whose RNA did not overlap with any gene body in GenecodeV29 in the sense orientation were classified as intergenic by tagtools and their STAR RNA alignments were extracted in a separate bam file. Only RNA reads with a STAR alignment score of *Q* = 255, a cognate DNA read with a Bowtie2 alignment score of *Q >* 15 were retained. The reads handling and filtering steps were performed as part of our ChAR-seq reads preprocessing Snakemake pipeline. These bam files were used as an input to StringTie2 with parameters `--fr --conservative -u -m 30 -p 4 -A` to produce one gtf file with de-novo gene models for each sample. The sample specific gtf files from the 2 ES and 2 DE ChAR-seq replicates were merged using StringTie2 with parameters `--merge -p 4 -m 30 -c 0 - F 0 -T 0` to produce a final a gtf file `intergenic.merged.gtf` with gene models for the UTL. This gtf file was used to generate a STAR index containing the gene models for the UTLs using command `STAR --runMode genomeGenerate --sjdbGTFfile intergenic.merged.gtf`. A dedicated Snakemake pipeline was run, similar to the full preprocessing pipeline described above, but starting from the tagtools step and using the UTL rather than the gencode gene models (and corresponding STAR indices) to produce pairs files corresponding to RNAs emanating from UTLs.

### Classification of UTLs

Each UTL was assigned 4 metrics or tags. i) We attributed each UTL a dominant Transposable Element (TE) family and a TE-score. For this task, we applied Classification of Ambivalent Sequences using K-mers (CASK)^85^ to the RNA-side of the ChAR-reads. CASK annotates each read with a candidate TE family (if any) based on its k-mer composition analyzed against a database of TE-specific k-mers built using the T2Tv1 genome assembly and T2T-CHM13 repeat annotations. Then, for each UTL, we identified the CASK annotation with the highest representation amongst all the reads (across the 2 ES and 2 DE replicates) mapped to this UTL. We assigned this annotation as the dominant TE family for this UTL and the proportion of reads from this UTL with this specific CASK annotation as its TE-score. ii) If the 5’ end of an UTL was within +/- 300 bp of a cis-regulatory element (CRE) active in either ES or DE cells, we annotated this UTL with the closest such CRE and its associated 7-group classification based on the Encode Registry of Regulatory Elements^67^ (file ID GRCh38-cCREs.bed). To determine active CRE in ES or DE cells, we selected, amongst the Encode Registry of Regulatory Elements (containing 1,063,878 human candidate CREs), those that overlapped with an ATAC-seq peak in that cell line. iii) UTLs whose 5’ end were within -200 bp to +100 bp of the 3’ end of a GencodeV29 gene body were flagged as candidate “readthrough.” iv) UTLs with at least 10% overlap with the antistrand of a GenecodeV29 gene body were flagged as candidate “antisense.” Finally, these 4 metrics and tags were combined to determine the final UTL classification using the following priority rule: i) UTLs with a dominant TE family of tRNAs and at TE-score greater than 10% were classified as tRNA-derived. ii) Remaining UTLs with a dominant TE family in {snRNA, snoRNA, scaRNA, srpRNA, scRNA, rRNA} and at TE-score greater than 10% were classified as snRNA-derived. iii) Remaining UTLs flagged as candidate readthroughs were classified as readthroughs. iv) Remaining UTLs with a CRE annotation in either ES or DE cells were classified as CRE-derived, and the subtype of CRE was selected from the ES cell annotation if the CRE was active in ES cells, and from DE cell annotation otherwise. v) Remaining UTLs with a TE-score greater than 50% were classified as repeat-derived, with the specific repeat family determined by their dominant TE family. vi) Remaining UTLs flagged as candidate antisense were classified as antisense. vii) All remaining UTL were classified as intergenic.

### Quantification of the RNA-DNA interactome dynamics

To compare the ChAR-seq RNA-DNA contact maps in ES versus DE cells, we repurposed the differential gene expression analysis tool DEseq2^94^. We applied DEseq2 in the interactome space (rather than the transcriptome space, as traditionally done in differential RNA-seq) using the number of ChAR-seq reads linking a specific RNA to a specific DNA locus, hereafter refer to an RNA-DNA interaction, as a separate rows in the input count matrix. We defined a DNA locus as either a 100 kb or 1 Mb genomic window (for Fig. 3), or a region surrounding the TSS of a protein coding gene as defined in the main text (for Fig. 6). The 4 ChAR-seq samples were included as columns of the count matrix. RNA-DNA interactions for which fewer than 2 samples had at least 10 reads were excluded from the count matrix and further analysis. The contact maps from exons, introns, and UTLs were analyzed in independent DESeq2 runs. The count matrices were generated in Python directly from the chartable objects that stored the contactome data. These matrices were imported in R and DESeq2 was run with all parameters set to their default values. Log2 Fold Change differential contacts maps shown in Fig. 3 were generated using the shrunken fold change estimates for each contact as returned by DESeq2. The apeglm method was used for shrinkage. This DESeq2 output was loaded into a chartable object in Python for computational handling and visualization tasks using chartools. Bar plots in Fig. 3b were produced using ggplot2 in R after converting the DESeq2 output into dplyr tibbles and applying appropriate transformations.

### Detection of RNA relocalization events during differentiation

Model 3 in Fig. 3c was tested by comparing the fold change between ES and DE cells for each RNA-DNA interaction with the fold change in total expression of the corresponding RNA in the caRNA transcriptome. To do so, we generated “expression only” contact maps, where the number of contacts between RNA *i* and genomic locus *j* was set equal to the total number of contacts made by RNA *i* in the observed map. For this analysis, genomic loci were defined using a 100 kb tiling partition of the genome. Because in these “expression only” maps, each row *i* (representing RNA *i*) is constant across the columns (representing the 100 kb-wide DNA loci), any information about the localization of individual RNAs is effectively removed and only the information about the abundance of each RNA is retained. We next applied DEseq2 in the interactome space as described above, but with the following modifications. First, the count matrix input to DEseq2 contained 8 samples/columns: the 2 ES and the 2 DE replicates of the observed contact maps and the 4 corresponding “expression only” maps. Second, we used a design matrix of the orm `∼cell + mapType + cell:mapType`, where the `mapType` covariate indicated whether the column corresponded to an observed ChAR-seq map or an “expression only” map, and the `cell` covariate indicated whether the column corresponded to a map in ES or DE cells. Third, the count matrix was prefiltered as above by removing interactions for which fewer than 2 samples had at least 10 reads, except that only the true observed samples (`mapType=observed`) were considered for the purpose of the filter. The interaction term `cell:rnaType` captured differences in the ES to DE dynamics in the true maps compared to the “expression only” maps. All interactions that had an FDR adjusted *p*-value associated with the `cell:rnaType` covariate smaller than 0.05 were flagged as “not explained by expression.” Maps shown in Fig. 3e and labeled as “Differential contacts explained by expression” were generated using the `apeglm` shrunken estimate of the regression coefficient associated with the `cell` covariate and with the reference level for `mapType` set to “expressionOnly.” This analysis was performed separately for maps corresponding to exons, introns, and UTLs.

### Computation of the *trans*- and *cis*-delocalization scores

For full details on the *trans*-delocalization scores please refer to Supplementary Note 2.

### trans-delocalization scores

Briefly, we defined the raw *trans*-de, l*_i_*ocal*_i_*ization*_i_* score*_i_* for*_i_* each R*_i_*NA as the ratio of the contact density of this RNA on *trans* chromosomes (number of contacts divided by the total length of the**_(1)_** *trans* chromosomes) over the contact density of this RNA on its *cis* chromosome. The raw delocalization score was difficult to interpret due to sample-specific biases and dependency in the chromosome of origin and expression (Supplementary Note 2, Supplementary Fig. 3). To regress out these biases and obtain a score that was comparable across RNAs and samples, we used a generalized linear model (GLM) and an empirical Bayes approach. First, we modeled the total number of *trans*-chromosomal contacts *N*_trans*,i*_ for each RNA *i* as independent Beta Binomial distributions. The Beta Binomial distribution accounts for both the sampling variation and the biological variation across RNAs, and was parameterized with the total number of reads *Ni* for RNA *i*, a mean *trans*-contact rate for RNA *i π_i_*, and an overdispersion parameter which we assumed constant across all RNA *γ*, such that

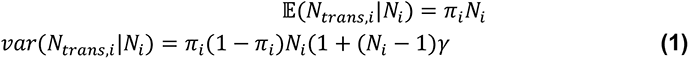

We captured the expression and chromosome biases by using a beta-binomial GLM and by including these effects as covariates in the GLM. Specifically, we used a logit link function for the mean *trans*-contact rate *pii* of the form

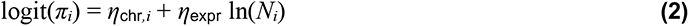

We next fit the Beta-binomial GLM using our ChAR-seq count data from mRNAs as a training set and conditioning on the total number of reads *Ni* for each RNA *i*. Fitting was performed using the `fit.gamlss` function from the `gamlss` package in R with the beta binomial family parameter and after loading the count data in a dplyr tibble and transforming the table appropriately for input into the fit function. RNAs with fewer than 50 total counts were removed and discarded from further analysis. Using the fitted beta-binomial GLM, we obtained for each RNA *i* an estimate for the mean *trans*-contact rate *π*_model*,i*_ and an associated Beta Binomial distribution with parameters *Ni*, *π*_model*,i*_ and *gamma model*, which we used as an Empirical Bayes prior. We performed a Bayesian update using the true observed number of *trans*-chromosomal contacts for RNA *i*, thereby obtaining a shrinkage estimate for the *trans*-contact rate*π*_post*,i*_. We defined the calibrated *trans*-delocalization score for RNA *i* Δ_trans, i_ as the log2 transformed ratio of the shrinkage estimate over the model prediction:

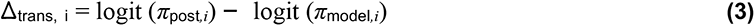

Delocalization scores were computed independently for each sample, and a final delocalization score for each RNA in each cell state was obtained by averaging the scores over the 2 replicates. For all delocalization score analyses, and only for these analyses, the pair files described in the above section (“ChAR-seq data processing and generation of pairs files”) were not used directly but further filtered to eliminate any possible remaining multi mappers on the RNA side, by using a more stringent multimapping threshold than STAR *Q* = 255. Specifically, the RNA side of the reads were realigned to hg38 using Bowtie2 and reads with *Q <* 40 were discarded from the pairs file for the delocalization score analysis.

### cis-delocalization scores

We defined the RNA travel distance *δ* for each ChAR-seq read corresponding to a *cis*- chromosomal contact as the distance between the mapping locus of the RNA and the mapping locus of the DNA. *cis*-delocalization scores were defined and computed similarly to the *trans*-delocalization scores, except for the following replacements: the number of *cis*-chromosomal contacts for RNA *i* was replaced with the number of contacts *Nδ<*1Mb*,i* such that the absolute RNA travel distance was smaller than 1 Mb, and the number of *trans*-chromosomal contacts was replaced with the number of contacts *Nδ>*1Mb*,i* such that the absolute RNA travel distance was greater than 1 Mb. The covariates for the GLM remained unchanged. Detection of RNAs with extreme delocalization scores The analysis described below was used for the *trans*- delocalization scores and was performed similarly for the *cis*-delocalization scores. Briefly, for each RNA and each sample, we computed the probability *p*_delocalized*,i*_ that a random sample drawn from the posterior distribution of the *trans*-contact rate *θ*post*,i* was larger than a random sample drawn from the GLM trained on the mRNA population. This probability was used as a *p*- value for identifying *trans*-delocalized RNAs. One *p*-value was obtained per RNA and per sample, and *p*-values from replicates were combined using Fisher’s method. Multiple hypothesis testing was corrected using the Benjamini Hochberg procedure. RNAs with an adjusted *p*-value smaller than 0.05 were declared as *trans*-delocalized. To identify RNAs on the other side of the distribution tail (ultra-localized RNAs) 1 *`−`p*_delocalized*,i*_ was used, and Fisher’s and BH methods were applied similarly. An RNA was declared ultra-localized if the resulting adjusted *p*-value was smaller than 0.05. All computations were performed in R. For further details please refer to Supplementary Note 3. Prediction of ChAR-seq contact maps using a generative model For mathematical details and a detailed discussion on the generative model, please refer to Supplementary Note 4. Briefly, the ChAR-seq dataset can be represented as a set of RNAs from an arbitrarily indexed transcriptome (i.e., RNA *i* refers to an RNA associated with the *i*^th^ gene in the transcriptome), and for each RNA *i*, a set of *N_i_* reads coming from this RNA whose RNA mapping coordinates are *`{`r_i,j_`}`j*=1*…N_i_* and DNA mapping coordinates are *`{`d_i,j_`}`j*=1*…N_i_* . We modeled for each RNA *i* the probability to observe any particular realization of the DNA mapping coordinates, conditional on knowing i) the set of RNA mapping coordinates and ii) the total number of contacts for this RNA on each chromosome. We modeled the *cis*- and *trans-chromosomal* contacts separately. For *cis*-contacts, we assumed the probability for an RNA emanating from coordinates *r* to contact locus *j* with coordinates *d_j_*, is proportional to: i) an RNA independent and DNA locus-dependent bias *b_j_* representing biological and technical variation of RNA localization and detection along the genome and ii) an interaction frequency dependent on the distance between the RNA and the DNA locus. The latter effect captures diffusion and tethering effects at short distances, whereby an RNA is more likely to interact with loci near its transcription site. Under this model, the probability to observe any specific localization pattern for RNA *i* in *cis* is given by a multinomial distribution of the form:

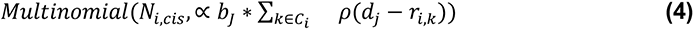

where *`C`i* is the set of indices amongst the reads from RNA *i*, for which the DNA-side maps to a locus in *cis*. For *trans*-contacts, we assumed that the probability for any RNA to contact locus *j* is only proportional to the DNA-bias. Under this model, the probability to observe any specific localization pattern for RNA *i* on a *trans* chromosome *c* is given by a multinomial distribution of the form

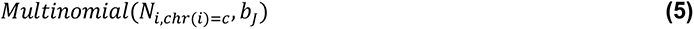

where *Ni,*chr(*i*)=*c* is the number of contacts made by RNA *i* on chromosome *c*. The DNA bias coefficients *b_j_* were estimated using the total coverage at each locus *j* from all the mRNAs originating from *trans* chromosomes. The distance dependent interaction frequency curve was estimated using the empirical distribution of RNA-DNA travel distance from all the protein coding RNAs. Maps shown across the manuscript and labeled as “model” were obtained by simulating a single realization of the *cis* and *trans* probabilistic models for each RNA in the transcriptome and for each target chromosome. Note that for each RNA, because of the conditional constraints, the total number of contacts on any specific chromosome are always equal in the simulated data and in the observed data. All simulations were performed in python as described in Supplementary Note 4, and the resulting maps were loaded in memory as `chartables` using `chartools` for analysis and plotting purposes.

### Detection of RNA-DNA contacts not predicted by the generative model

To compare the true observed ChAR-seq RNA-DNA contact maps to those predicted by the generative model, we applied DEseq2 in the interactome space as described in the section “Quantification of the RNA-DNA interactome dynamics” with the following modifications. First, the count matrix input to DEseq2 contained 8 samples/columns : the 2 ES and the 2 DE replicates of the true “observed” contact maps, and the 4 corresponding “model” maps, obtained by a single simulation of the generative model. Second, the design matrix was set to `∼ cell + observedORmodel + cell:observedORmodel`, where the `observedORmodel` covariate indicated whether the column corresponded to “observed” or “model” ChAR-seq map. Third, the count matrix was prefiltered by removing interactions for which fewer than 2 samples amongst the “observed” samples had at least 10 reads. The interaction term `cell:rnaType` captured differences between the observed and modeled data that were specific to either ES to DE cells. All interactions whose `apeglm` shrunken estimate of the regression coefficient associated with the `observedORmodel` covariate was greater than *log*2(1.3) and had an FDR adjusted *p*-value smaller than 0.0 5 were flagged as “not explained by model.” We computed the regression coefficient associated with the `observedORmodel` and its *p*-value in ES and DE cells separately, by setting the reference level for the `cell` covariate to ES and DE, respectively, before running DEseq2. This analysis was performed separately for maps corresponding to exons, introns, and UTLs. Maps shown in Fig. 5d-h, and Fig. 6b labeled as “model’ or “mod” were generated using the `apeglm` shrunken estimate of the regression coefficient associated with the `observedORmodel` covariate. The DESeq2 outputs were loaded into a chartable object in Python using chartools for visualization tasks. Bar and line plots in Fig. 5e were produced using ggplot2 in R after converting the DESeq2 output into dplyr tibbles and applying appropriate transformations.

### External data used in this study

Hi-C data in Fig. 5 were loaded in HiGlass from the Krietenstein et al. 2019 (H1 hESCs) dataset^95^, visualized at 2kb resolution after ICE normalization, and manually aligned with the ChAR-seq, ATAC-seq, H3K27ac and H3K4me3 tracks plotted in IGV based on their genomic coordinates. H3K27ac and K3K4me3 tracks in Fig. 2b and Fig. 5g were generated using ChIP-seq data in H7 hESCs cells and H7 cells differentiated into definitive endoderm from GSE127202^45^. PolII localization peaks used for the metagene analysis in Extended Data Fig. 5e were obtained by running MACS2 on H9 ChIP-seq data from GSE105028^96^.

### Data and code availability

All ChAR-seq, RNA-seq and ATAC-seq sequencing data generated as part of this study are available by request and will be made publicly available on GEO upon publication. Packages released as part of this study and Snakemake pipelines used to preprocess the ChAR-seq and RNA-seq data are available on github at the specific repositories described above. All data analysis code and code to generate the figures are available by request and will be available on github upon publication. Any additional information required to reanalyze the data reported in this paper is available from the lead contact upon request.

## Acknowledgements

This research was supported by the National Institutes of Health (NIH) grant NIH R01 HG009909 to WJG and AFS. OKS was supported by the Training Grant NIH T32-GM113854-02 and NSF-GRFP, and by the Stanford Center for Systems Biology Seed Grant Award. Some of the computing for this project was performed on the Sherlock cluster. We would like to thank Stanford University and the Stanford Research Computing Center for providing computational resources and support that contributed to these research results. We thank Viviana Risca for assistance with the protocols and helpful discussions, and Ali Shariati for assistance with cell culture. We thank the members of the Straight lab for advice and feedback on the manuscript.

## Contributions

CL, OKS, DJ, WJG and AFS conceptualized the study. OKS, DJ, and KAF performed cell culture work. DJ prepared RNA-seq and ATAC-seq libraries, and processed the ATAC-seq data. CL, OKS and DJ performed the ChAR-seq experiments and library preparation. CL developed the concepts related to RNA localization patterns, developed and wrote the computational tools to analyze the data, and performed the computational and statistical analysis. OKS, DJ and KAF helped with data analysis. CL performed visualizations. KAF helped with figures. CL wrote the original draft of the manuscript. CL, OKS, DJ, KAF, WJG and AFS reviewed and edited the manuscript. WJG and AFS provided resources and acquired funding. AFS supervised the study.

