## Supplementary Information for "Global mapping of RNA-chromatin contacts reveals a proximity-dominated connectivity model for ncRNA-gene interactions"

**Content:**

- Extended Data Figures 1-6
- Supplementary Note 1: Computational pipeline for ChAR-seq reads preprocessing
- Supplementary Note 2: Delocalization scores
- Supplementary Note 3: Identification of RNAs with extreme delocalization scores
- Supplementary Note 4: Modeling the ChAR-seq contact maps
- Supplementary Tables (attached files)

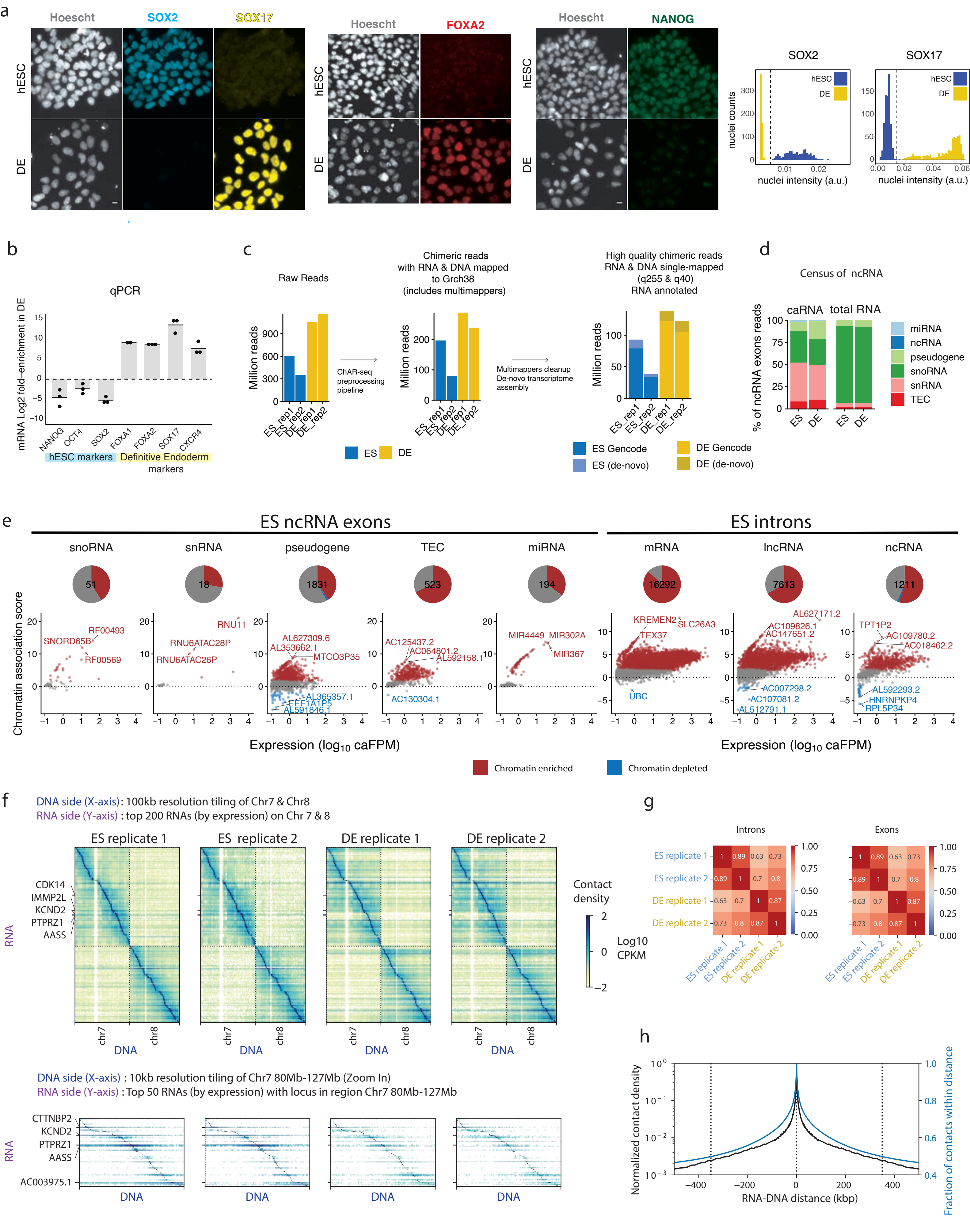

a
Hoescht
SOX2
SOX17
FOXA2
NANOG
hESC
DE
SOX2
SOX17
nuclei counts
nuclei intensity (a.u.)
b
qPCR
mRNA Log2 fold-enrichment in DE
NANOG
OCT4
SOX2
FOXA1
FOXA2
SOX17
CXCR4
hESC markers
Definitive Endoderm markers
c
Raw Reads
Chimeric reads with RNA & DNA mapped to Grch38 (includes multimappers)
High quality chimeric reads RNA & DNA single-mapped (q255 & q40) RNA annotated
Million reads
ChAR-seq preprocessing pipeline
Multimappers cleanup De-novo transcriptome assembly
ES_rep1
ES_rep2
DE_rep1
DE_rep2
ES
DE
ES Gencode
ES (de-novo)
DE Gencode
DE (de-novo)
d
Census of ncRNA
caRNA
total RNA
% of ncRNA exons reads
miRNA
ncRNA
pseudogene
snoRNA
snRNA
TEC
e
ES ncRNA exons
ES introns
snoRNA
snRNA
pseudogene
TEC
miRNA
mRNA
lncRNA
ncRNA
51
18
1831
523
194
16292
7613
1211
Chromatin association score
SNORD65B
RF00493
RF00569
RNU11
RNU6ATAC28P
RNU6ATAC26P
AL627309.6
AL353662.1
MTCO3P35
AL365357.1
EEF1A1P5
AL591846.1
AC125437.2
AC064801.2
AL592158.1
AC130304.1
MIR4449
MIR302A
MIR367
KREMEN2
SLC26A3
TEX37
UBC
AL627171.2
AC109826.1
AC147651.2
AC007298.2
AC107081.2
AL512791.1
TPT1P2
AC109780.2
AC018462.2
AL592293.2
HNRNPKP4
RPL5P34
Expression (log10 caFPM)
Chromatin enriched
Chromatin depleted
f
DNA side (X-axis): 100kb resolution tiling of Chr7 & Chr8
RNA side (Y-axis) : top 200 RNAs (by expression) on Chr 7 & 8
ES replicate 1
ES replicate 2
DE replicate 1
DE replicate 2
CDK14
IMMP2L
KCND2
PTPRZ1
AASS
RNA
DNA
chr7
chr8
Contact density
Log10 CPKM
DNA side (X-axis) : 10kb resolution tiling of Chr7 80Mb-127Mb (Zoom In)
RNA side (Y-axis) : Top 50 RNAs (by expression) with locus in region Chr7 80Mb-127Mb
CTTNBP2
AC003975.1
g
Introns
Exons
h
Normalized contact density
Fraction of contacts within distance
RNA-DNA distance (kbp)

**Extended Data Fig. 1. Validation of H9 hESCs differentiation into definitive endoderm, ChAR-seq reads statistics, chromatin enrichment details, and replicability of the ChAR-seq maps, related to Fig. 1. a**, Validation of H9 hESCs differentiation into definitive endoderm by widefield immunofluorescence. H9 cells are stained against Sox2, Sox17, FoxA2 and Nanog, pre- and post-differentiation. Bar plots show quantification of Sox2 and Sox17 staining. **b**, Validation of H9 hESCs differentiation by qPCR of key pluripotency state marker genes (OCT4, SOX2, NANOG) and definitive endoderm marker genes (FOXA1, FOXA2, SOX17, CXCR4) in ES and DE cells. qPCR results shown as relative expression levels calculated using the $2^{\Delta\Delta CT}$ method with the PBGD housekeeping gene for normalization. Data points represent 3 technical replicates. **c**, Number of raw reads obtained for each cell type and replicate (left), number of reads left after quality filter for which both the RNA and DNA mapped to the genome (including multimappers, no Q score filtering, middle panel), and final number of chimeric reads with high confidence alignment and annotation (Bowtie2 alignment Q score >=40 on the DNA side, STAR alignment score 255 on the RNA side, single gene annotation, either from Gencode v29 or from the de-novo transcriptome as described Fig. 2). These high quality chimeric reads were used for all the analysis in this paper, except where indicated otherwise. **d**, Percentage of ncRNA reads originating from specific subtypes of ncRNAs. The ncRNA subtypes are obtained from Gencode v29 and further simplified as indicated in Table S7. **e**, Left group: Breakdown of the chromatin-association score versus expression scatter plot shown in Fig. 1d (ncRNA, right) by subtypes of ncRNAs in ES cells. Right group: same plots as in Fig. 1d but for introns of individual RNAs rather than exons. Data are displayed as in Fig. 1d. **f**, Replicability of the ChAR-seq maps. ChAR-seq maps for individual replicates in ES and DE cells are shown at two different resolutions as indicated. All the maps are shown as introns and exons of individual genes summed together. Color indicates contact per genomic kb per million reads (CPKM). **g**, Cross-correlation of the 100kb resolution maps across cell lines and replicates, and separately for RNAs originating from exons, introns, and UTL. The cross-correlation for a pair of maps is computed as the Pearson correlation of the pixel intensities in the maps (i.e., the contact rates between an RNA and a 100kb genomic interval). Maps with all chromosomes and all RNAs expressed above 0.1 FPM in the caRNA transcriptome are used to compute the correlations. **h**, Distance-dependent interaction curve, showing the likelihood for an mRNA exon to contact a genomic site as a function of the RNA travel distance, defined as the distance between the RNA transcription locus (mapping coordinate of the RNA-derived side of the corresponding ChAR-seq read) and the target DNA locus (mapping coordinate of the DNA-derived side of the ChAR-seq read).

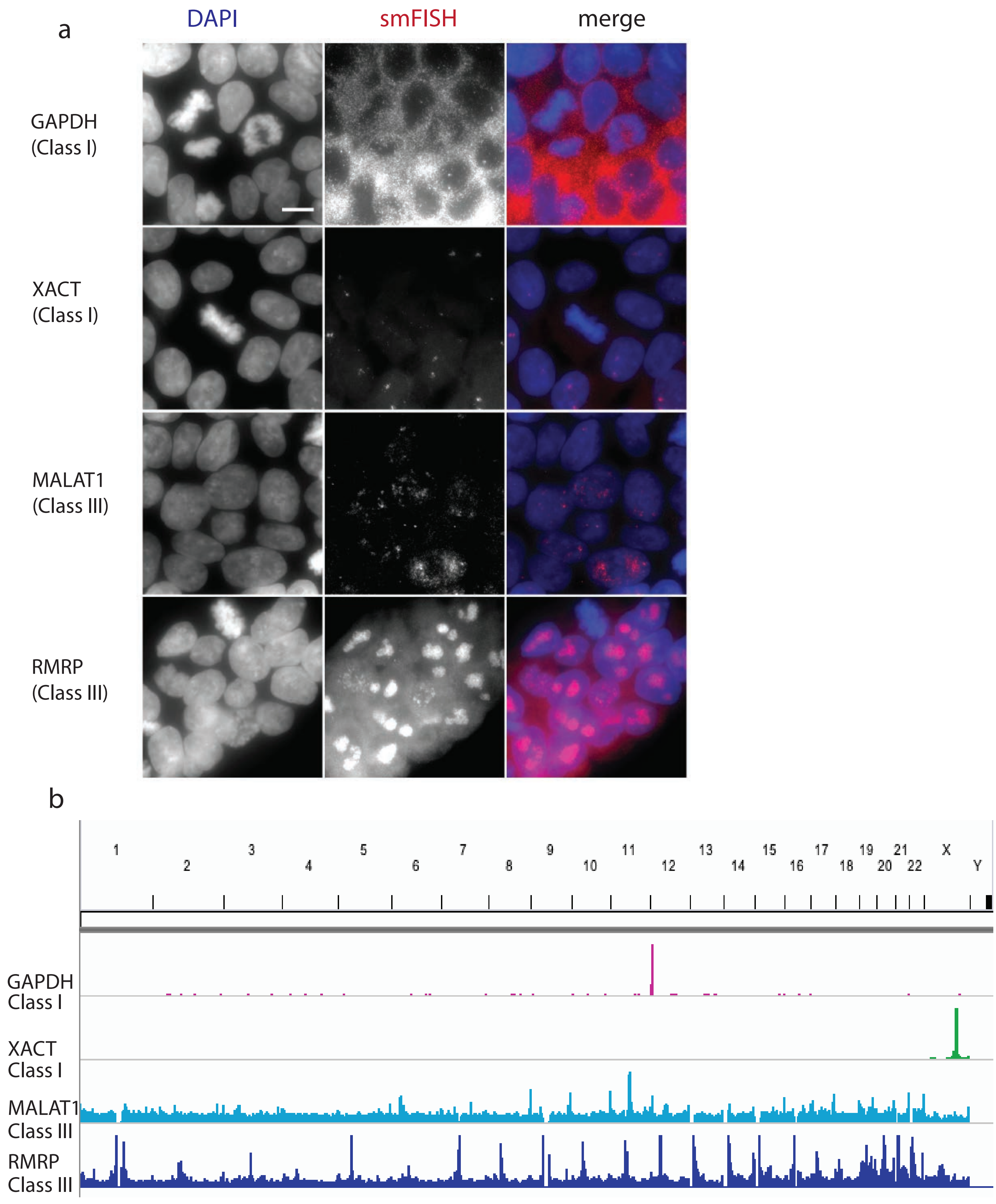
a
DAPI
smFISH
merge
GAPDH
(Class I)
XACT
(Class I)
MALAT1
(Class III)
RMRP
(Class III)
b
X
Y
GAPDH
Class I
XACT
Class I
MALAT1
Class III
RMRP
Class III

**Extended Data Fig. 2. smFISH validation of the localization patterns of select RNAs, related to Fig. 1. a**, Representative localization patterns of GAPDH, XACT, MALAT1, and RMRP in individual cells, as measured by smiFISH(Tsanov et al. 2016). **b**, Chromatin localization profiles determined by ChAR-seq in ES cells of the RNAs shown in a.

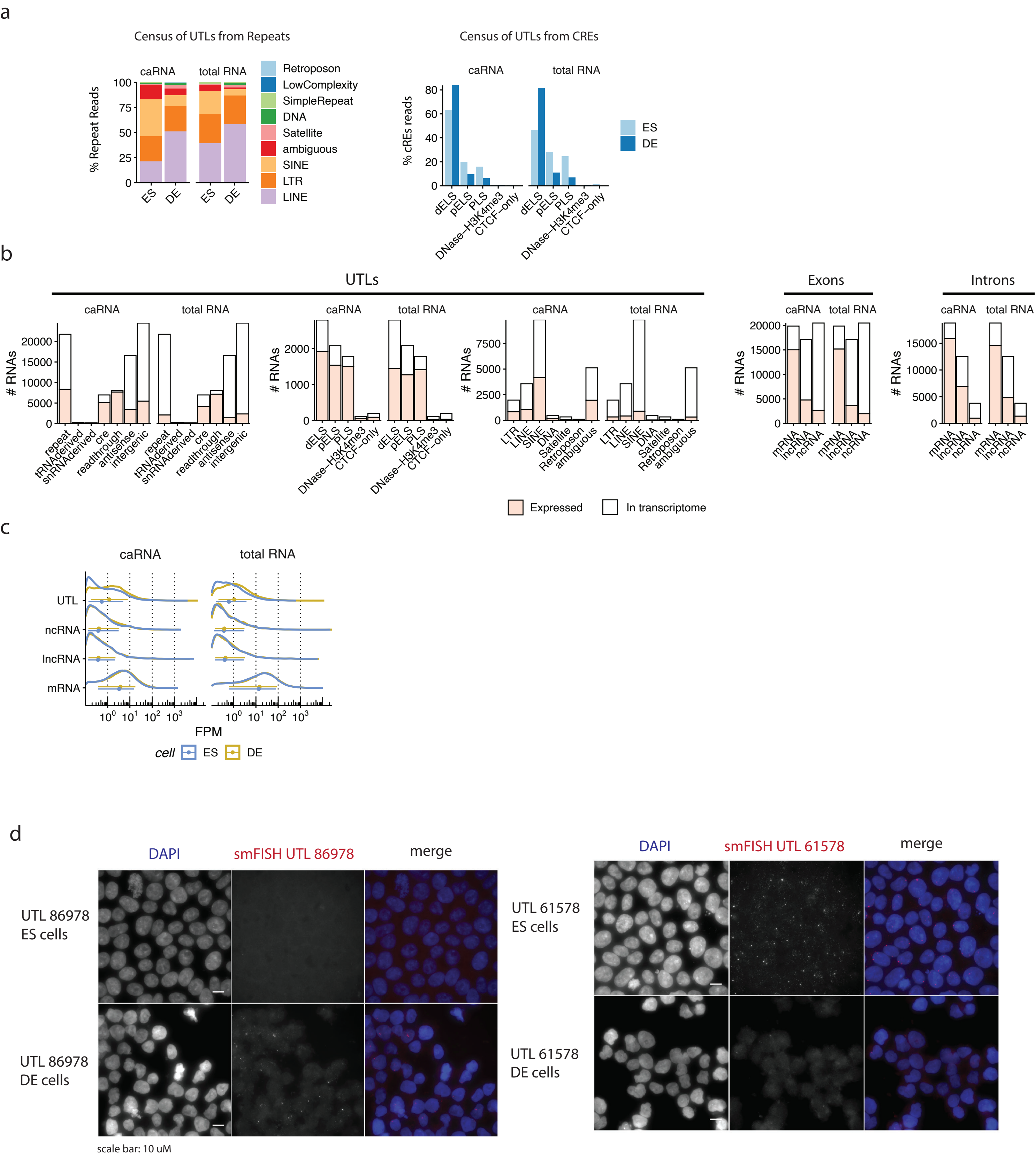
a
Census of UTLs from Repeats
Census of UTLs from CREs
caRNA
total RNA
% Repeat Reads
Retroposon
LowComplexity
SimpleRepeat
DNA
Satellite
ambiguous
SINE
LTR
LINE
ES
DE
% cREs reads
dELS
pELS
PLS
DNase–H3K4me3
CTCF–only
b
UTLs
Exons
Introns
### RNAs
repeat
tRNAderived
snRNAderived
cre
readthrough
antisense
intergenic
mRNA
lncRNA
ncRNA
Expressed
In transcriptome
c
UTL
FPM
cell
d
DAPI
smFISH UTL 86978
merge
UTL 86978 ES cells
UTL 86978 DE cells
scale bar: 10 uM
smFISH UTL 61578
UTL 61578 ES cells
UTL 61578 DE cells

**Extended Data Fig. 3. Detailed composition of the unannotated transcribed loci (UTLs), related to Fig. 2. a**, Percent of UTLs reads originating from each subtype of repeat (left, relative to reads annotated as repeat-derived), and each subtype of Cis-regulatory elements (right, relative to reads annotated as CRE-derived). Cis-regulatory elements are classified using the 7-group classification from the Encode Registry of Regulatory Elements(ENCODE Project Consortium et al. 2020) (pELS=proximal Enhancer Like Sequence, dELS=distal Enhancer Like Sequence, PLS = Promoter Like Sequence, see STAR Methods). **b**, Diversity of the UTLs. Bar plots show the absolute number of RNAs expressed at FPM above 0.1 for each Gencode type of exons and introns, and for each type of UTL. The FPM value refers to the maximum of the ES and DE FPM values. “In transcriptome” values refer to the total number of RNA of each type in either GencodeV29 or in the catalog of UTLs generated in this study (Table S3) **c**, Distribution of the expression level of the UTLs compared to exons of annotated mRNAs, lncRNAs and ncRNAs. **d**, Subcellular localization, determined by single molecule RNA-FISH, in ES and DE cells of two cell-state specific UTLs: UTL 86978 (left panels) and UTL 61578 (right panels). DNA staining is shown in the first column, RNA-FISH signal in the second column, and a merge in the third column. ES cells are in the top row and DE cells in the bottom row. Scale bar = 10µm.

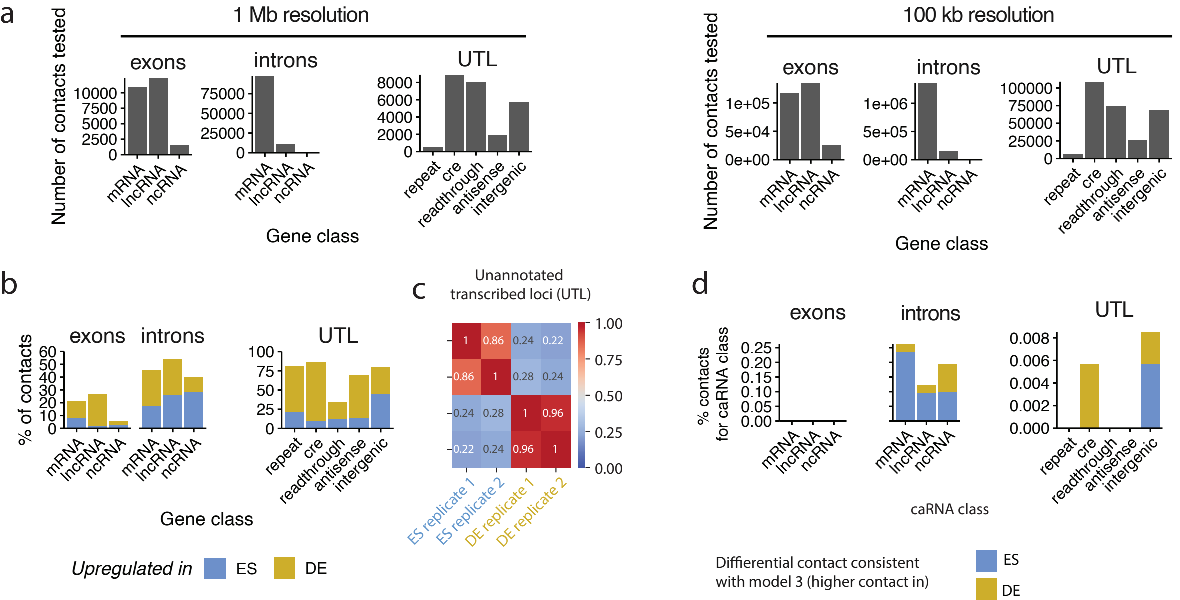
a
1 Mb resolution
100 kb resolution
Number of contacts tested
exons
introns
UTL
mRNA
lncRNA
ncRNA
repeat
cre
readthrough
antisense
intergenic
Gene class
b
% of contacts
Upregulated in
ES
DE
c
Unannotated transcribed loci (UTL)
ES replicate 1
ES replicate 2
DE replicate 1
DE replicate 2
d
% contacts for caRNA class
caRNA class
Diffierential contact consistent with model 3 (higher contact in)

**Extended Data Fig. 4. RNA-DNA contacts of UTLs are more dynamic during differentiation than those of annotated RNAs, related to Fig. 3. a**, Number of RNA-DNA interactions tested for differential representation in ES versus DE cells at 1 Mb DNA resolution (left) and 100 kb DNA resolution (right), by class of RNA. **b**, Quantification by RNA class of the percentage of interactions upregulated in DE or ES cells amongst all interactions tested in that class (interactions with >10 counts in at least one replicate in ES or DE), at 1 Mb DNA resolution. Same analysis as in Fig. 3b, but at 1 Mb resolution on the DNA side rather than 100 kb resolution. **c**, Cross-correlation of the RNA-DNA contacts maps of UTLs at 100kb resolution. Correlations are computed as in Extended Data Fig. 1f. **d**, Percentage of differential interaction not explained by differential RNA expression, at 100 kb resolution, relative to the total number of interactions tested within the RNA class.

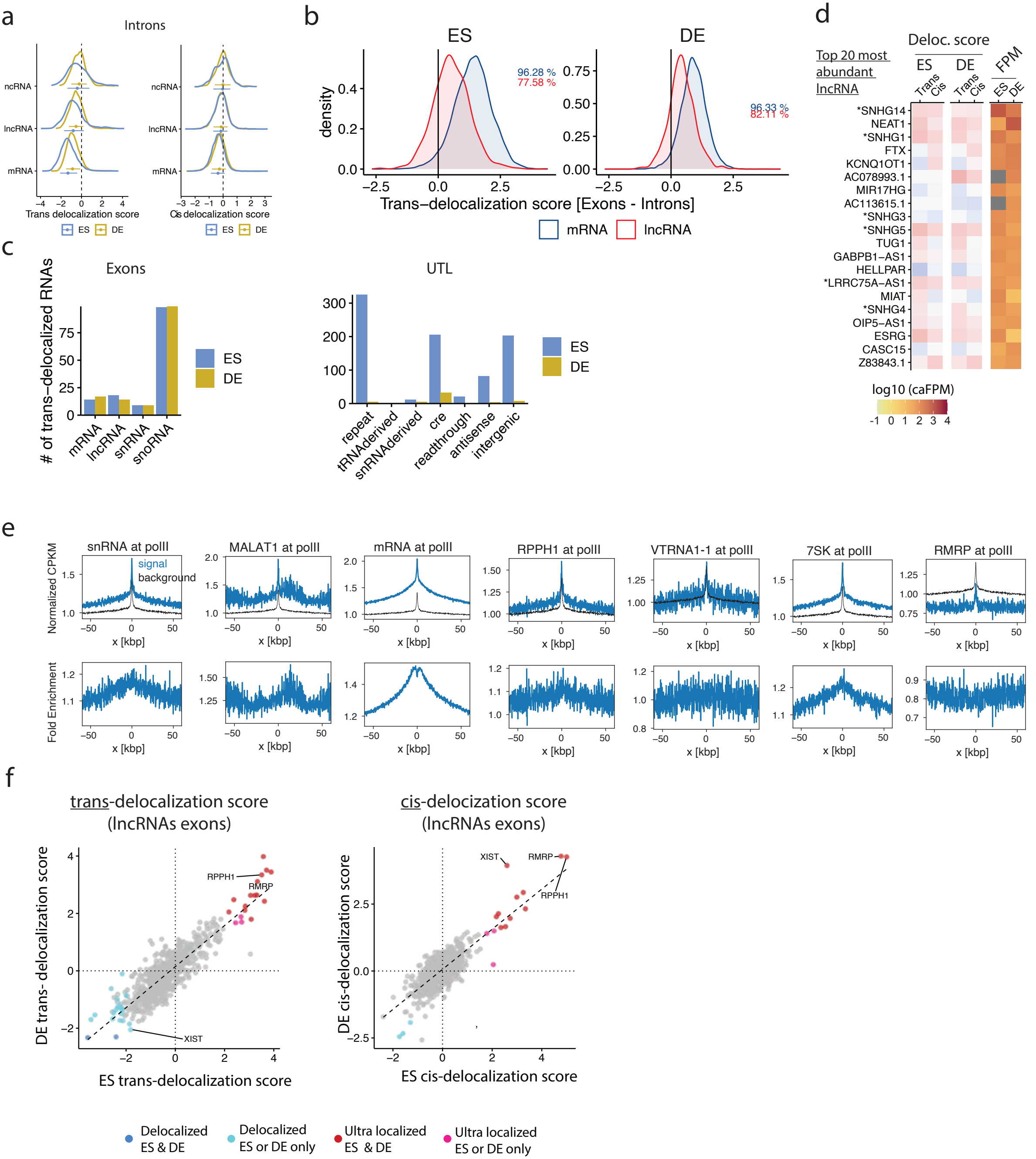

a
Introns
ncRNA
lncRNA
mRNA
Trans delocalization score
Cis delocalization score
ES
DE
b
ES
DE
density
96.28 %
77.58 %
96.33 %
82.11 %
Trans−delocalization score [Exons - Introns]
mRNA
lncRNA
c
Exons
UTL
### of trans−delocalized RNAs
mRNA
lncRNA
snRNA
snoRNA
repeat
tRNAderived
snRNAderived
cre
readthrough
antisense
intergenic
ES
DE
d
Top 20 most abundant lncRNA
Deloc. score
ES
DE
Trans
Cis
FPM
*SNHG14
NEAT1
*SNHG1
FTX
KCNQ1OT1
AC078993.1
MIR17HG
AC113615.1
*SNHG3
*SNHG5
TUG1
GABPB1−AS1
HELLPAR
*LRRC75A−AS1
MIAT
*SNHG4
OIP5−AS1
ESRG
CASC15
Z83843.1
log10 (caFPM)
e
snRNA at polII
MALAT1 at polII
mRNA at polII
RPPH1 at polII
VTRNA1-1 at polII
7SK at polII
RMRP at polII
signal
background
Normalized CPKM
Fold Enrichment
x [kbp]
f
trans-delocalization score (lncRNAs exons)
cis-delocization score (lncRNAs exons)
DE trans- delocalization score
ES trans-delocalization score
DE cis-delocalization score
ES cis-delocalization score
RPPH1
RMRP
XIST
Delocalized ES & DE
Delocalized ES or DE only
Ultra localized ES & DE
Ultra localized ES or DE only

**Extended Data Fig. 5. Additional data on delocalization scores and broadly localized RNAs, related to Fig. 4. a**, Distribution of trans- (left panel) and cis- (right panel) delocalization scores for introns of mRNAs, lncRNAs, and ncRNAs. Bars below each distribution represent median and interquartile range. **b**, Distribution of the difference in trans-delocalization score between the exons and introns of the same RNA and as function of the type of RNA (mRNA or lncRNA) and cell type. Percentages indicate the percentage of RNAs for which the exonic trans-delocalization score is larger than the intronic trans-delocalization score. **c**, Absolute number of RNAs in each category classified as delocalized (either cis- or trans-localized at FDR 0.05). **d**, Heat maps showing the cis and trans delocalization scores and chromatin abundance in ES and DE cells for the 20 most abundant lncRNAs (excluding those identified as cis- or trans-delocalized). **e**, Metagene plots showing the levels of several RNA categories (mRNAs, snRNAs) and individual RNAs (7SK, MALAT1, VTRNA1-1, RMRP) near genomic loci with PolII ChIP-seq peaks. PolII peaks were obtained from GSE105028(Lyu et al. 2018). For each panel, the top plot shows the pileup of the specific RNA or RNA group (blue line), and the pileup for the background RNAs, defined as the total signal of all mRNAs localized on trans-chromosomes, which captures DpnII site density bias, accessibility bias, and other forms of non-specific localization bias (Methods and Supplementary Note 4). Each signal is displayed as a contact density (CPKM = contacts per 1k genomic bp per 1 million reads), normalized by the median metagene background contact density in 10kb bins in a 1Mb region around the feature, (so that the background decays to 1.0 far from the feature center). The bottom plot shows the RNA fold enrichment over background. f, Scatterplots showing the trans- (left) and cis-delocalization scores (right) for individual lncRNAs in DE versus ES cells. Black lines show linear regression output.

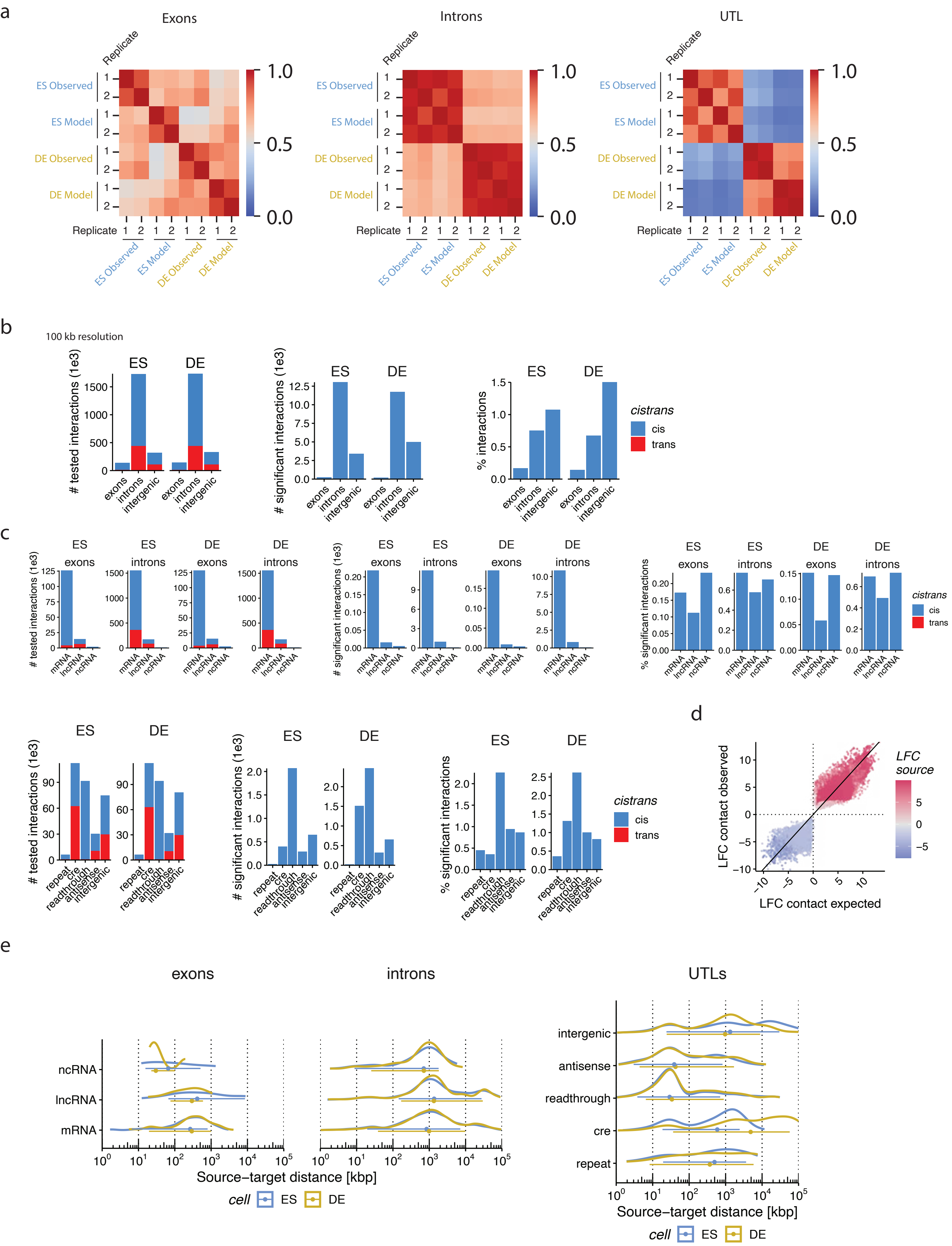
a
Exons
Introns
UTL
Replicate
ES Observed
ES Model
DE Observed
DE Model
1.0
0.5
0.0
b
100 kb resolution
ES
DE
### tested interactions (1e3)
### significant interactions (1e3)
% interactions
exons
introns
intergenic
cistrans
cis
trans
c
mRNA
lncRNA
ncRNA
% significant interactions
repeat
cre
readthrough
antisense
intergenic
d
LFC contact observed
LFC contact expected
LFC source
e
exons
introns
UTLs
Source–target distance [kbp]
cell
ES
DE

**Extended Data Fig. 6. Expression-distance model predicts >99% of RNA-chromatin contacts, related to Fig. 5. a**, Cross-correlation of the 100 kb resolution RNA-DNA contact maps obtained from the true ChAR-seq data and predicted by the generative model. Cross-correlation matrices are shown separately for the RNAs from introns, exons, and for UTLs. The cross-correlation for a pair of maps is computed as in Extended Data Fig. 4b. **b**, Number of interactions tested for enrichment over model (left), number of interactions identified as over-represented in the observed data compared to the model (significant interactions, middle), and proportion of significant interactions in relation to the total number of tested interactions in each RNA category (right). For this analysis, interactions are defined at 100 kb resolution on the DNA side. Cis (resp. trans) indicate interactions where the RNA and DNA are on the same (resp. on a different) chromosome. **c**, Same quantification as in b, but broken up by RNA type. **d**, Scatter plot showing for each lncRNA-gene contact its observed log2 fold change during ES to DE cells differentiation (LFC) by ChAR-seq, and its predicted LFC by the generative model. Contacts are defined as in Fig. 7a-b. Color indicates the LFC of the lncRNA level on chromatin (shrunken estimate computed using DESeq2 applied to the RNA-side of the ChARseq data, Table S1). **e**, Distribution of the RNA-DNA travel distance for interactions significantly above the model. Related to Fig. 5f but broken up by type of RNA.

#### Supplementary Note 1 : Computational pipeline for ChAR-seq reads preprocessing

Demultiplexed fastq files were preprocessed using a custom snakemake(Mölder et al. 2021) pipeline (https://github.com/straightlab/charseq-pipelines**).** The main goal of the pipeline is to produce pairs files containing information about each RNA-DNA contact. Pairs files are tab separated files with the first 4 columns describing the RNA and DNA coordinates of each RNA(cDNA)-DNA chimeric read and other relevant annotations for each contact stored in subsequent columns. These files are in pairix compatible format, so they can be indexed to enable efficient 2D queries using pairix. The pipeline produces intermediates files such as split fastq files corresponding to the RNA and DNA side of each read, bam files for the RNA and DNA alignments, etc. The pipeline steps are described below. A summary of the pipeline workflow is depicted in Supplementary Fig. 1. Read statistics at various steps of the pipeline for the 4 ChAR-seq samples are shown in Supplementary Fig. 2.

##### Deduplication, adapters trimming and debridging

PCR duplicates were removed using clumpify.sh from BBMap v38.84 (parameters dedupe=t subs=0 reorder=f). Reads were quality thresholded and sequencing adapters were trimmed using Trimmomatic v0.38(Bolger et al. 2014) (parameters: PE ILLUMINACLIP:<trimfasta>:2:30:12 SLIDINGWINDOW:10:10 MINLEN:61, with <trimfasta> pointing to the definition file for the adapters). To detect chimeric cDNA-DNA reads where the ChAR-seq bridge may span across read mates 1 and 2 in the paired end data, read pairs were merged using Pear v0.9.6(Zhang et al. 2014) (parameters: -p 0.01 -v 20 -n 50). Three fastq files were generated: one containing reads whose mates were successfully merged, and a pair of files containing paired end reads that could not be merged. Merged (M) and unmerged (U) reads were separately processed as single end and paired end reads, respectively. M reads were scanned to detect the ChAR-seq bridge sequence, and reads containing a single occurrence of that sequence were “debridged,” i.e., they were split into a rna.fastq and dna.fastq file corresponding to the sequences of the RNA (cDNA) and DNA side of the chimeric molecules, respectively. Bridge sequence detection and debridging were performed using Chartools v0.1, a custom package to process ChAR-seq data released as part of this study **(https://github.com/straightlab/chartools-dev).** Specifically, we used the Julia(Bezanson et al. 2017) script debridge.jl with the -s option (single end). Reads that did not contain the bridge sequence or contained multiple occurrences of the bridge were dumped into separate fastq files and discarded from further analysis. U reads were debridged similarly except using the paired-end mode of the debridge.jl script (without -s option). In that case, only reads where the bridge sequence was found a single time across both mates (i.e., in either read 1 or read 2 but not both) were kept for subsequent analysis. For these read pairs, the mate that did not contain the bridge was discarded, and the mate containing the bridge was split into a rna.fastq and dna.fastq file. In these split fastq files from the M and U branches, reads were either left unchanged or reverse complemented depending on the orientation of the charseq-bridge, in such a way that i) all RNA reads in rna.fastq are represented in their sense orientation (the right most nucleotide in the sequence corresponds to where the 3’ end of the RNA where it was ligated to the bridge) and ii) all DNA reads in dna.fastq are in sense orientation with respect to the bridge (the left most nucleotide in the sequence corresponds to the 5’ end of the DNA where it ligated to the bridge). This operation is part of the debridge.jl script and transparent to the user. Finally, rna.fastq files and dna.fastq files from the U and M processing branches were merged, to obtain final single-ended rna.fastq and dna.fastq files, with the read IDs matching line by line across these two files.

##### Length filtering and removal of ribosomal RNA reads

ChAR-seq reads whose RNA- or DNA-derived sequence were shorter than 15bp were filtered out using the custom chartools script filter_short_reads_pairs.sh, to produce length filtered rna.fasq and dna.fastq files with the read IDs matching line by line across these two files). Reads whose RNA-derived sequence mapped to a ribosomal RNA were removed by aligning the rna.fastq file to a fasta file of ribosomal sequences downloaded from NCBI using Bowtie2(Langmead and Salzberg 2012) (parameters: -q --very-sensitive --norc). Reads with one or more valid alignments were filtered out of rna.fastq using picard. The corresponding DNA-derived sequence of these reads were filtered out of dna.fastq using the chartools script decon_reads_pairs.sh.

##### Alignment and annotation with known genes

DNA reads were aligned against hg38 using Bowtie2 v2.3.4.1 (parameters: --ultra-sensitive), producing a dna.bam file. RNA reads were aligned against hg38 using STAR(Dobin et al. 2013) and a gtf annotations file obtained from Gencode V29 (parameters: --outFilterMultimapNmax 10 --outSAMmultNmax 10 --outSAMattributes All --outReadsUnmapped None --outSAMunmapped Within --outMultimapperOrder Random --quantTranscriptomeBan Singleend). To assign RNA reads to specific genes, we used tagtools (https://github.com/straightlab/tagtools, a package released as part of this study which annotates STAR aligned reads with the set of genomic features they overlap with, amongst a user defined set of reference features. We applied tagtools using the transcript definition gtf file from GencodeV29, producing a bam file rna.exons.bam with all the reads (referred to as “exonic” reads in this study) that fully overlapped with known transcripts, and containing a supplementary field AN:<transcriptID> field indicating the most likely transcript of origin. Reads that did not fully overlap with known transcripts were selected from the original rna.fastq file based on their read ID using Picard, and realigned with STAR using the same transcript definition gtf file but with an index produced with --sjdbGTFtagExonParentTranscript ID --sjdbGTFfeatureExon gene parameters. This allowed us to obtain reads that aligned to intronic regions of gene bodies. These reads were annotated using tagtools to produce a bam file (rna.introns.bam) containing a supplementary AN:<geneID> field indicating the most likely gene of origin. Finally, reads that did not fully overlap with known gene bodies but were not classified as having “too many” mapped loci or as unmapped were separated into a third bam file (rna.intergenic.bam), which was used later with StringTie2(Kovaka et al. 2019) to detect novel transcriptional units and generate the unannotated transcribed loci (UTL) catalog.

##### Pairs file generation

The aligned DNA reads in dna.bam and the aligned and annotated RNA reads were combined read-by-read into a pair file using the pairup function in chartools. Separate pair files were generated for reads whose RNA was annotated in the STAR/tagtools step as exonic, intronic, and intergenic reads. Each pair file contains the mapping coordinates of the DNA and of the RNA, the most likely transcript and gene of origin identified with tagtools, and other information about the alignments, such as the alignment quality score or the number of gene annotations compatible with the RNA mapping locus. These pairs files are in a pairix compatible format and were indexed using pairix(Lee et al. 2022).

##### Final Filtering

Pairs files were filtered to remove multimapping reads and reads with low mapping scores on either the RNA (STAR Q<255) or DNA (Bowtie2 Q<40) side. Using tagtools-derived annotation of the RNA reads, we also removed reads for which the RNA could not be either attributed to an unambiguous gene defined in Gencode or to a single intergenic or antisense locus. Finally, we discarded reads whose RNA overlapped with a region on the ENCODE blacklist(Amemiya et al. 2019). The filtered pairs files were used for all the analysis in this work.

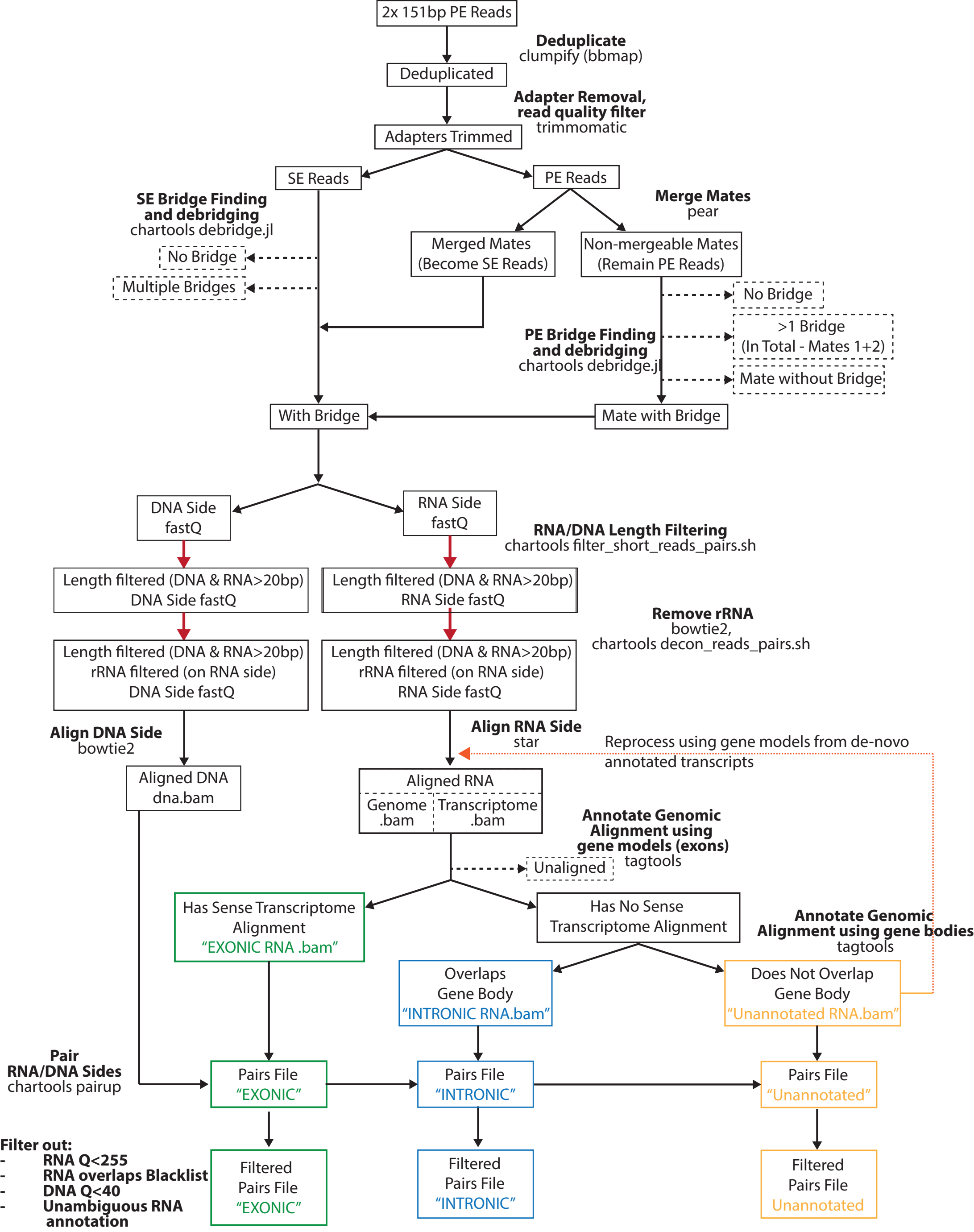

Red arrows indicates RNA & DNA side processed together as a pair (to maintain readID match line by line on RNA and DNA)
Dotted orange arrow indicates add-on to pipeline to annotate reads that don't overlap with Gencode Genes after de-novo transcriptome assembly (STAR methods)

**Supplementary Figure 1. Diagram of the ChAR-seq preprocessing computational pipeline, showing the processing steps and associated tools.** The pipeline takes the fastq files from the sequenced ChAR-seq libraries as input, and outputs “pairs” files which essentially contain the RNA and DNA coordinates of each contact, along with an annotation of the genes from which the RNA originates.

#### Supplementary Note 2: Delocalization scores

##### Modeling the contact rate on *trans* chromosomes

We denote by RNA $i$ the $i^{th}$ RNA in an arbitrarily indexed transcriptome. Let $N_{cis,i}$ be the number of *cis*-chromosomal contacts for RNA $i$ (contacts with a locus on the RNA chromosome of origin), and $N_{trans,i}$ be the number *trans*-chromosomal contacts (contacts with a locus on a chromosome other than the RNA chromosome of origin). We define the raw fraction of contacts in *trans* $\theta_{raw,i} = N_{trans,i}/(N_{trans,i} + N_{cis,i})$, and the raw *trans*-delocalization score $\Delta_{trans,\ raw,\ i}$ as

$$\Delta_{trans, raw, i} = log\left(\frac{\theta_{raw,i}}{\left(1-\theta_{raw,i}\right)} * \frac{L_{cis}}{L_{trans}}\right) \qquad \textbf{(S1)}$$

where $L_{cis}$ and $L_{trans}$ are the total length of the *cis* chromosome and of all the *trans* chromosomes combined, respectively. The raw *trans*-delocalization is effectively, in log scale, the ratio of the contact density (i.e. number of contacts per unit genomic length in the target chromosomes) in *trans* over *cis*. We noted that, when looking at the distribution of the raw delocalization scores across mRNAs, these distributions were shifted in location across replicates, indicating the presence of sample specific biases (Data S2 [Figure 1b]). Furthermore, the raw *trans*-delocalization scores were also correlated with expression, and surprisingly, we noted that RNAs from exons and introns behaved differently: the raw delocalization score of an exonic RNA was positively correlated with its abundance, while the delocalization score of an intronic RNA was negatively correlated with its abundance (Supplementary Fig. 3). To regress out these biases and obtain calibrated delocalization scores that are comparable across samples and RNAs, we modeled the number of contacts in *trans* using a Generalized Linear Model (GLM) with a beta-binomial response. In absence of biological noise, since each contact can be either in *trans* or in *cis*, it is reasonable to assume that, conditional on the total number of reads $N_i = N_{trans,i} + N_{cis,i}$, the number of *trans* contacts follows a binomial distribution with unknown success probability $\theta_i$. Distributions with constrained mean-variance relationship such as the binomial or Poisson distribution typically do not work well with sequencing data due to the presence of unmodelled biological or technical variation. Thus, we modeled $\theta_i$ as a Beta distribution, such that, conditional on $N_i$

$$N_{trans,i}|\theta_i, N_i \sim Binomial\left(N_i, \theta_i\right)$$

$$\theta_i \sim Beta\left(\pi_i \frac{1-\gamma}{\gamma}, (1 - \pi_i)\frac{1-\gamma}{\gamma}\right) \qquad \textbf{(S2)}$$

The resulting compound distribution for $N_{trans,i}$ is a Beta Binomial. The parametrization of the Beta distribution was chosen such that the $\pi_i$ is the mean *trans*-contact rate and $\gamma$ is the overdispersion parameter such that $E(N_{trans,i} \mid N_i) = \pi_i N_i$ and $var(N_{trans,i} \mid N_i) = \pi_i(1 - \pi_i)N_i (1 +(N_i - 1)\gamma)$. This approach is motivated by the similarity of the problem with that of estimating methylation rate at CpGs site from bisulfate sequencing data, where the available data are the number of methylated and unmethylated reads, and for which a Beta binomial model has been proposed(Dolzhenko and Smith 2014; Park et al. 2014) [13,14].

We assumed that the overdispersion parameter is constant across RNAs, and that the mean *trans* contact rate $\pi_i$ is only function of the chromosome of origin and of the total level of RNA $i$ on chromatin. Specifically, we used the a logit link function and a regression model of the form:

$$logit\left(\pi_i\right) = \eta_{chr,i} + \eta_{expr} ln\left(N_i\right) \qquad \textbf{(S3)}$$

where $logit(x) = 1/(1 - x)$.

Since mRNAs are not expected to have any defined chromatin targets, we reasoned that the number of *trans* contacts for mRNAs should most closely follow the model. Thus we used mRNAs to fit the regression coefficients $\eta_{chr,i}$ and $\eta_{expr}$, and the overdispersion parameter $\gamma$. Fit was performed using the gamlss package in R with the Beta Binomial family(Stasinopoulos and Rigby 2007).

To estimate the *trans*-contact probability $\pi_i$ for individual RNA *i*, we used an empirical Bayes method. Specifically, we used the observed counts $N_{cis,i}$ and $N_{trans,i}$ to compute the posterior estimates for the beta distribution parameters $\pi_{post,i}$ and $\gamma_{post}$ . Using the prior estimates given by the fitted model $\pi_{model,i}$ = invlogit ($\eta_{chr,i} + \eta_{expr} \ln(N_{cis,i} + N_{trans,i})$) and $\gamma = \gamma_{model}$, the posterior estimates for the beta distribution parameters are obtained using the following update formula:

$$\alpha_{post,i} = \alpha_{model,i} + N_{trans,i}$$

$$\beta_{post,i} = \beta_{model,i} + N_{cis,i} \qquad \textbf{(S4)}$$

where invlogit is the inverse logit function and the parameters $\alpha$ and $\beta$ are the canonical parameters of the beta distribution related to the desired contact probability and overdispersion parameter by $\pi = \frac{\alpha}{\alpha+\beta}$ and $\gamma = \frac{1}{\alpha+\beta+1}$. Using the empirical Bayes estimator for the contact probability, we obtain a shrinkage estimator for the delocalization score, simply as $logit(\pi_{post,i}) + log(L_{cis}/L_{trans})$. Finally, we define the calibrated *trans*-delocalization score as the difference between the shrinkage estimator and the fitted model :

$$\Delta_{trans,i} = logit(\pi_{post,i}) - logit(\pi_{model,i}) \qquad \text{(S5)}$$

The calibrated *trans*-delocalization score $\Delta_{trans,\ i}$ is effectively a log fold difference in the *trans*- vs *cis*-contact density ratio for RNA *i* compared to *trans*- vs *cis*-contact density ratio for an "average" mRNA of the same expression level.

#### Modeling the contact rate far for the transcription locus and *Cis*-delocalization scores

For each RNA-DNA contact (each RNA-DNA read in the raw ChAR-seq data), we denote by $\delta$ the distance between the mapping locus of the RNA and the mapping locus of the DNA. We hereafter refer to $\delta$ as the RNA "travel" distance (see Data S3 [Technical Note] for further details regarding this distance). We denote by $N_{far,i}$ the number of contacts from RNA *i* for which the absolute travel distance is larger than *w*, and by $N_{close,i}$ the number of contacts for which the absolute travel distance is smaller than *w*. Throughout this study we used *w* = 10 Mb as the threshold for considering a contact as close (proximal *cis* interaction) versus far (distal *cis* interaction). We defined the raw *cis*-delocalization score (at threshold *w*) for RNA *i* similarly to its *trans*-delocalization score, but replacing $N_{trans,i}$ by $N_{far,i}$ and $N_{cis,i}$ by $N_{close,i}$. We also replaced the genomic space normalization factors $L_{trans}$ and $L_{cis}$ by $L_{cis} - w$ and *w* respectively. We used a similar GLM with a beta binomial response to model the number of counts $N_{far,i}$ conditional on the total number of counts in *cis* $N_{cis,i} = N_{close,i} + N_{far,i}$, and trained this model on the population of mRNAs. Following the same approaches as in our *trans*-score analysis, we finally obtained a calibrated *cis*-delocalization scores $\Delta_{cis,i}$.

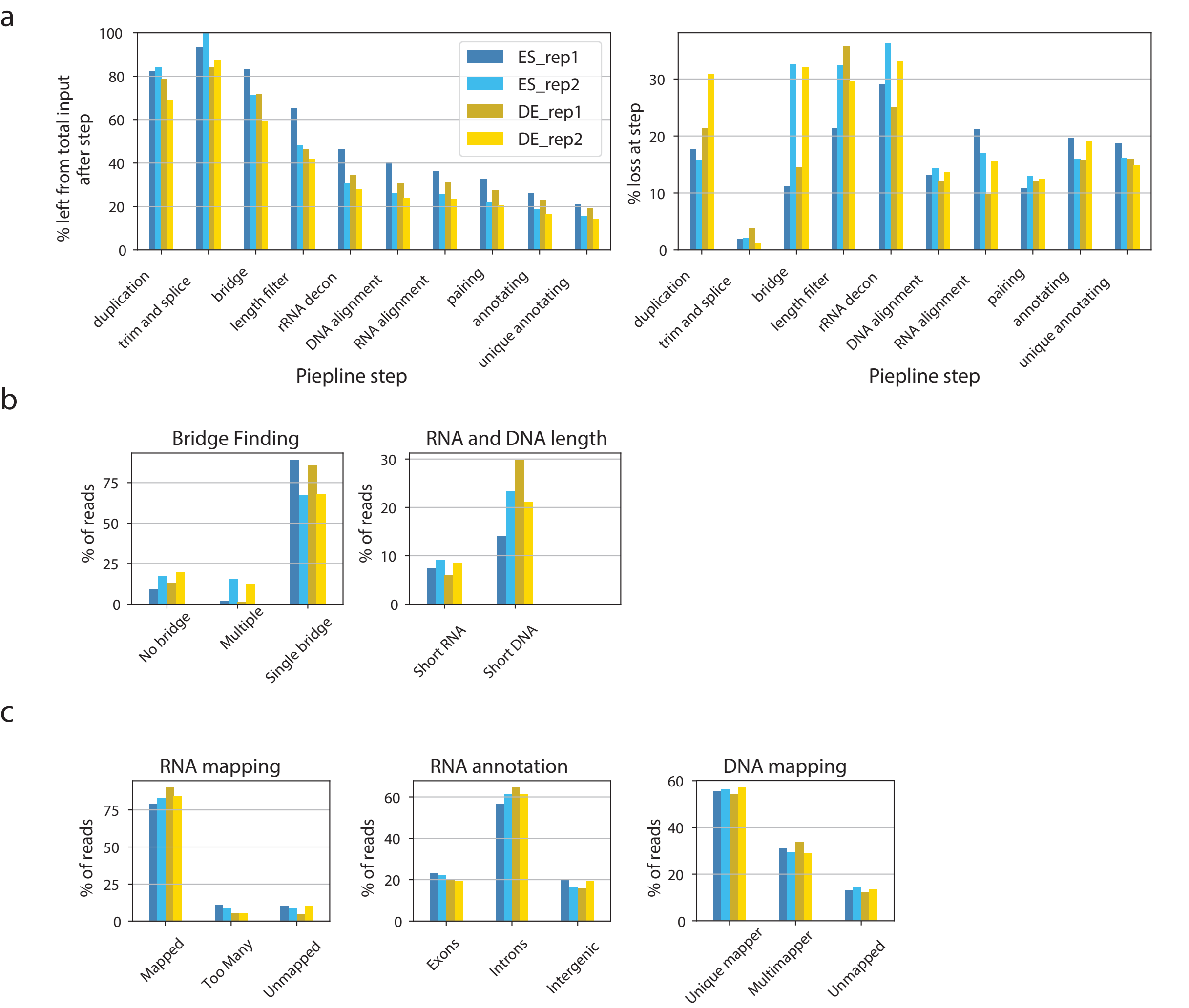

a
% left from total input after step
Piepline step
ES_rep1
ES_rep2
DE_rep1
DE_rep2
% loss at step
duplication
trim and splice
bridge
length filter
rRNA decon
DNA alignment
RNA alignment
pairing
annotating
unique annotating
b
Bridge Finding
% of reads
No bridge
Multiple
Single bridge
RNA and DNA length
Short RNA
Short DNA
c
RNA mapping
Mapped
Too Many
Unmapped
RNA annotation
Exons
Introns
Intergenic
DNA mapping
Unique mapper
Multimapper
Unmapped

**Supplementary Figure 2. ChAR-seq libraries QC: reads statistics at various stages of the preprocessing pipeline. a**, Loss of reads not meeting quality filters during the preprocessing pipeline. Left panel shows the percentage of reads left after each stage of the processing pipeline, relative to the initial number of raw reads. Right panel shows the fraction of reads lost at each stage of the pipeline, relative to the number of reads left after the prior stage. **b**, Details on the bridge finding and length filtering, showing the breakdown of the reasons why reads were filtered out at this stage. For the bridge finding step, “None” indicates no bridge sequence was found in the raw read, and “Multiple” indicates several occurrences of the bridge sequence were found in the raw read. Only reads with a single occurrence (“Single”) of the bridge sequence were kept. For the length filtering step, short RNA (resp. short DNA) indicates that the RNA (resp. DNA) side of the read was shorter than 15bp. **c**, Details on the alignment step, showing the percentage of reads that were discarded during the alignment of either the RNA or DNA side, due to too many mapped locations or no mapping location (as reported by STAR and Bowtie2, respectively). Percentages shown are relative to the number of reads left after the rRNA decontamination step. Reads that passed the alignment filtering step on both the RNA and DNA side were selected for the pairing stage. Reads that passed only the RNA or only the DNA alignment filter or none of them were discarded at the pairing stage. The RNA annotation bar plot shows the percentage of reads relative to the number of reads passing the RNA alignment stage as described above, which overlapped exclusively to exons, or to introns (including exons-introns junctions) defined in Gencode V29. Intergenic indicates reads that were not contained within a gene body or were antisense to a gene.

### Supplementary Note 3: Identification of RNAs with extreme delocalization scores

#### *trans*-delocalization

To label RNAs with extreme *trans*-delocalization scores, we computed the probability that a random sample drawn from the posterior distribution of $\pi_i$ is larger than a random sample drawn from the prior distribution obtained by training the GLM on the mRNA population. This probablity was computed using the analytical formula defined in(Miller 2015).This probability is the right-tail probability of a compound model where a Beta distribution (with overdispersion $\gamma_{\text{post},i}$) is sampled after sampling its mean according another Beta distribution with mean $\pi_{\text{prior},i}$ and overdispersion $\gamma_{\text{prior}}$. Thus this probability can be interpreted as the *p*-value, which we denote by $p_{\text{delocalized},i}$, for the *trans*-contact rate being positive and more extreme than that of an mRNA. We obtained one such *p*-value per RNA and per condition and replicate. RNAs with fewer than 50 counts (across all its DNA targets) were excluded from the analysis. To combine the *p*-values from different replicates, we used Fisher's method and obtained a final *p*-value per RNA in ES cell $p_{\text{delocalized},i,\text{ES}}$ and one final *p*-value per RNA in DE cells $p_{\text{delocalized},i,\text{DE}}$. Finally, we adjusted these *p*-values for multiple hypothesis testing using the Benjamini Hochberg procedure(Stasinopoulos and Rigby 2007) (independent adjustment for the set of ES and DE *p*-values, number of tests equal to the number of RNA tested in the corresponding condition). We declared an RNA as *trans*-delocalized in ES or DE when the its corresponding adjusted *p*-value was smaller than 0.05. Similarly, we used $p_{\text{ultra-localized},i} = 1 - p_{\text{delocalized},i}$ as the *p*-value for the *trans*-contact rate being negative and more extreme than that of an mRNA, used the Fisher Method to combine replicates, and adjusted these *p*-values with the Benjamini Hochberg procedure. We declared an RNA as ultra-localized (with respect to its *trans* contacts) when its adjusted *p*-value was smaller than 0.05.

*Trans*-delocalization scores and associated *p*-values for each RNA are given in Table S8 before combining replicates, and Table S4 after combining replicates.

#### *cis*-delocalization

Following a similar procedure with the *cis*-delocalization scores, we obtained for each RNA *i*, and an adjusted p-value that allows us to identify RNAs with extreme *cis*-delocalization scores. As RNA with an adjusted p-value smaller than 0.05 was labeled as *cis*-delocalized (when its delocalization scores were positive), or ultra-localized (with respect to its *cis* contact patter, when its delocalization scores were negative). These data are given in Table S9 before combining replicates, and Table S4 after combining replicates.

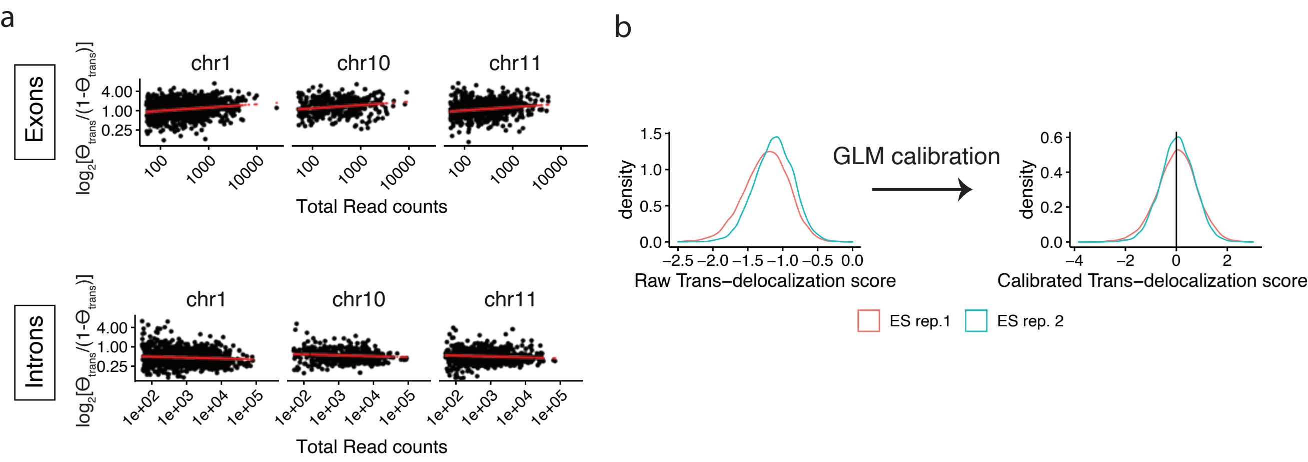
a
Exons
Introns
chr1
chr10
chr11
$\log_2[\Theta_{trans}/(1-\Theta_{trans})]$
4.00
1.00
0.25
100
1000
10000
1e+02
1e+03
1e+04
1e+05
Total Read counts
b
GLM calibration
density
Raw Trans−delocalization score
Calibrated Trans−delocalization score
ES rep.1
ES rep. 2

**Supplementary Figure 3. Calibration of trans-delocalization scores using a Generalized Linear Model a**, Uncalibrated trans-delocalization score for individual mRNAs exons (top) or introns (bottom) as a function of their expression and chromosome of origin (chr1, chr10, and chr11 shown as subpanels), and GLM fit (red line). GLM is defined in equation S3 **b**, Distribution of the raw trans-delocalization scores of mRNAs for the two ChARseq replicates in ES cells as defined in equation S1, and of the calibrated trans-delocalization scores as defined in S5. The raw trans-delocalization scores show sample biases which are regressed out after calibration.

#### Supplementary Note 4: Modeling the ChAR-seq contact maps

##### Definition of the generative model

We denote by $M$ the RNA-DNA contact matrix, where $M_{i,j}$ is the raw number of contacts for RNA $i$ at DNA locus $j$. More specifically, RNA $i$ refers to the $i^{th}$ RNA in an arbitrarily indexed transcriptome as above, and DNA locus $j$ refers to the $j^{th}$ interval in an arbitrary set of genomic intervals (typically the $j^{th}$ tile in a tiling partition of the genome). We also label the chromosome of locus $j$, chr($j$), and its mean genomic coordinate, $d_j$. Our goal is to predict $M$ using a simple generative model in such a way that deviations from the prediction carry meaningful biological information. The ChAR-seq sequencing experiment can be described as a random sampling process, where the probability to draw a contact between RNA $i$ and locus $j$ is, in absence of any bias, proportional to i) the true level of RNA $i$ on chromatin and ii) the true probability that RNA $i$ physically interacts with locus $j$, as opposed to with any other particular locus. However, both of these assumptions do not hold true due to the presence of technical and biological biases. A standard approach used to analyze Hi-C data is to correct for these biases by applying a matrix balancing operation, such as the Vanilla Coverage (VC), Knight-Ruiz (KR) or Iterative Correction (ICE) normalization(Imakaev et al. 2012; Knight and Ruiz 2012). Here, we do not use these approach for two reasons. First, rather than balancing the raw data, we seek to derive a generative model that reflects the probability to observe the contact matrix $M$ , including the effects due to the biases. Second, in contrast to Hi-C data, the matrix $M$ is not symmetric and different biases, hereafter referred to as RNA-side and DNA-side biases, affect the rows and columns of $M$ , such that the standard Hi-C balancing algorithms are not directly applicable. RNA-side biases may originate from differences in RNA-bridge ligation efficiency across RNA species, protection by RNA-binding proteins or RNA structure, and RNA mappability. DNA-side biases may stem from accessibility and mappability differences across genomic loci and from sites with non-specific affinity for RNAs.

In the scope of this paper, we are interested in examining the binding patterns of individual RNAs across the genome, not in quantifying the relative abundance of RNA species at individual loci. Thus, we do not need to model the full process giving rise to $M$ . Rather, we assign ourselves a simpler objective, which is to model the process giving rise to $M$ conditional on observing the RNA-side of the data. This greatly simplifies the problem as we can discard the RNA biases. More rigorously, denoting $N_i$, the total number of observed ChAR-seq reads from RNA $i$, we condition our model on fixing the mapping coordinates (3' end coordinates) $\{r_{i,k}\}_{k=1...N_i}$ of the RNA-side of these reads to those observed in the data. In this conditional setting, the probability to observe a read mapping on the RNA-side to $r_{i,k}$ and on theDNA-side to locus $j$ is proportional to i) the true probability $\Pi_r i,k,j$ that a physical fragment of RNA $i$ with a 3' end coordinate $r_{i,k}$ interacts with genomic locus $j$ and ii) a locus specific DNA-bias coefficient $b_j$ capturing all of the various DNA-side biases described above. Note that because a gene may span over several hundreds of kbp, the loci $r_{i,k}$ can be far from each other. Thus, it is not accurate to approximate $r_{i,k}$ by a single coordinate such as the TSS or TES of the gene. To further refine the probability model, we need at this point to separately examine the case of *cis*-chromosomal and *trans*-chromosomal contacts. For *cis*-chromosomal contacts, we reasoned that in the absence of directed interactions, RNA-chromatin contacts arise as a result of the diffusion of the RNA. Diffusion starts from the RNA transcription locus. A contact may occur with locus $j$ only if the RNA is within some capture radius of the locus for the ligation reactions during the library preparation. Thus, the true contact probability $\Pi_r i,k,j$ should be, at minimum, function of i) an unknown probability $\Lambda_{i\rightarrow \mathrm{chr}(i)}$ that the RNA remains on its chromosome of origin (denoted by chr($i$)), rather than diffuses to a different chromosome, and ii) the genomic distance between the RNA and DNA loci. Thus we can write, for all loci $j$ on cis-chromosome:

$$\Pi_{r_{i,k},j} \propto \Lambda_{i\to chr(i)} \times \rho_{chr(i)}\left(\epsilon_i * \left(d_j - r_{i,k}\right)\right) \quad \textbf{(S6)}$$

where $\rho chr(i)(\delta)$ is an unknown distance-dependent RNA-DNA interaction frequency which we allow to be distinct on distinct chromosomes, and $\epsilon_i$ indicates the orientation of the gene (+1 if the RNA is on the + strand and −1 otherwise). $\delta$ denotes the distance between the mapping locus of the RNA-side and DNA-side of a contact read, defined earlier as the RNA "travel" distance. The sign of $\delta$ indicates whether the DNA target is located downstream ($\delta > 0$), or upstream ($\delta < 0$) of the RNA locus, in reference to the transcription direction. Thus $\epsilon_i$ corrects for the orientation of the gene.

For *trans*-chromosomal contacts, we propose the simple model where the true contact probability for a locus on a *trans*-chromosome *C* is uniform across this chromosome and is proportional only to the probability that RNA *i* diffuses to *C* rather than to any other chromosome. Thus, for all loci *j* on *trans*-chromosome *C*:

$$\Pi_{r_{i,k},j} \propto \Lambda_{i\to C} \quad \textbf{(S7)}$$

The coefficient $\Lambda_{i\to C}$, which we refer to as the inter-chromosomal transfer coefficient, models the overall contact rate for RNA *i* to chromosome *C*. Consequently, this coefficient does not affect the shape of the contact profile, only the relative global levels of the RNA on different chromosomes. For the purpose of detecting local anomalies in the observed contact patterns, such as peaks at discrete loci, we can essentially leave the inter-chromosomal transfer coefficient out of the model parameters. Specifically, we simplified our model by fixing the marginals corresponding to the total number of contacts for RNA *i* on each chromosome *C*, and setting them to the observed number of contact. Let us denote $n_{i,C}$ the number of contacts in the observed data made by RNA *i* on chromosome *C*. Combining all of the reads from the same RNA *i* together, we obtain a generative model giving the distribution of RNA *i* on each chromosome as a multinomial distribution:

$$M_{i,j} \sim \begin{cases} Multinomial\left(n_{i,chr(j)}, \alpha_i * b_j * \sum_{k\in C_i} \rho(d_j - r_{i,k})\right) & \text{if } j \text{ in } cis \\ Multinomial\left(n_{i,chr(j)},\right) & \text{if } j \text{ in } trans \end{cases} \quad \textbf{(S8)}$$

where chr(*j*) indicates the chromosome on which the target locus *j* is located, and $C_i$ is the set of indices amongst the reads from RNA *i*, for which the DNA-side maps to a locus in *cis* (simply speaking, we use the $r_{i,k}$ only for the reads that map to a *cis* RNA-DNA contact). In this expression the first argument of the multinomial is the number of draws, and the second argument is the events probability vector. $\alpha_i$ is a normalization coefficient such that the sum of the entries in the events probability vector equals to 1. Note that $b_j$ is already normalized on each *trans*-chromosome, as discussed below.

Alternatively, if one wishes to capture broad chromosome scale features, such as excess of an RNA across a specific chromosome, we can combine the generative model with the GLM model used to calculate the *trans*-delocalization scores. Specifically, we can fix the marginals so that the total number of contacts on each chromosome *C* is equal to that predicted by the GLM. This simply amounts to replacing, in the equation above, $n_{i,C}$, with its GLM prediction, $\hat{n}_{i,C}$. This approach, which we term “generative model with trans-contact rate prediction” was used to generate the model tacks in Figure 4f.

**Interpretation of the generative model**

It is clear that the simple generative model described above cannot, in principle, capture all of the complexities of the data for 3 main reasons. First, diffusion of an RNA occurs in 3D, so the genomic distance $\delta$ used in the distance-dependent RNA-DNA interaction curve does not accurately represent the physical proximity between the two cognate loci, especially in the context of the complex 3D conformation of the genome. Even for *trans*-chromosomal interactions, specific arrangements of the chromosomes in 3D may bring diverse *trans*-chromosomal loci in close proximity to the RNA transcription locus, invalidating the assumption that the true contact probability is uniform across a *trans* chromosome. Second, the diffusivity of an RNA is likely dependent on its genomic context (e.g., whether it is within a heterochromatin 290 or euchromatin region), its secondary structure (e.g., smaller RNAs may be able to diffuse further awayfrom their locus), and its affinity for specific nuclear factors. Third, by design of the model, it does not encode any potential "affinity driven" interaction between a specific RNA and a specific target locus(Engreitz et al. 2016). However, this is not a limitation but a feature, as deviations from the model indicate biologically interesting interactions, namely those that cannot be explained by 1D proximity and expression.

**Estimation of the DNA-bias**

We reasoned that because most mRNAs are unlikely to have any site-specific chromatin activity, their contact patterns on chromatin should be non-specific and most closely follow the generative model. Furthermore, although some of the model assumptions may be invalid for some individual RNAs as discussed above, we reasoned that deviations from the model are likely to average out when the contact patterns of all mRNAs are aggregated. Thus, we empirically estimated the DNA-bias vector $b_j$ by counting, for each locus $j$, the total number of contacts at this locus from all mRNAs originating from *trans*-chromosomes (hereafter referred to as *trans*-mRNAs). Let $U_j$ be the set of indexes $i$ such that RNA $i$ is a *trans*-mRNA at locus $j$, we computed the DNA-bias vector as:

$$b_j = \sum_{i \in U_j} M_{i,j} \qquad \textbf{(S9)}$$

We then normalized this vector such that the partial sum of its entries across each individual chromosome is equal to 1:

$$b_j \rightarrow b_j / \sum_{k|chr(k)=chr(j)} b_k \qquad \textbf{(S10)}$$

$b_j$ can be interpreted as a distant-independent and RNA-independent DNA-bias. Estimation of the distance-dependent RNA-DNA interaction frequency for *cis*-chromosomal interactions Based on the argument previously mentioned that most mRNAs are unlikely have any site-specific interactions, we also used mRNAs to estimate the distance-dependent RNA-DNA interaction frequency $\rho_C(\delta)$ for each chromosome $C$. We estimated the interaction frequency at logarithmically spaced positive distances $\{\delta_j\}_{j=0 \ldots n}$ , and negative distances $\{\delta_j = -\delta_{-j}\}_{j=-n \ldots -1}$, where $\delta_0 = 0$ and $\delta_j = 1 \ldots n_\delta \delta\ \delta$ spanned 10 bp to 100 Mbp (an arbitrarily large value larger than the size of the largest chromosome). To do so, we tallied across all mRNAs transcribed on chromosome $C$, the number of *cis* contacts $O_{C,\delta j}$ with a travel distance within $[\delta_j, \delta_{j+1})_{j=(-n\ )\ldots(n\ -1)}$. To account for the edge effects (the fact that RNAs$\delta\ \delta$ cannot make contacts at distances beyond their distance to the chromosome edge), we divided $O_{C,\delta j}$ by the maximum number of contacts that could be observed at a given distances $A_{C,\delta j}$ if all RNAs were forced to localize at this distance $\delta_j$ when possible. We finally normalized the resulting vector by the sum of its entries to obtain an estimate of the distance-dependent RNA-DNA interaction frequency:

$$\rho_C(\delta_j) = \frac{O_{C,\delta_j} / A_{C,\delta_j}}{\sum_{j=-n\delta}^{n_\delta - 1} O_{C,\delta_j} / A_{C,\delta_j}} \qquad \textbf{(S11)}$$

***Generation of predicted contact map.***

$$\Pi_j \propto b_j |\{k | d_{sim,i,k} = d_j\}| \quad \textbf{(S12)}$$

where $|S|$ denotes the cardinality of the set $S$. Note that this probability is zero at all the loci that were not sampled during the first step, making this simulation very efficient (typically the number of reads from an RNA is much smaller than the number of target loci $j$, especially when the target loci are high resolution ($\lessapprox 1Mb$) tiling partitions of the genome).We simulated the *trans*-localization patterns in a manner similar to the *cis*-localization patterns, but skipped the travel distance sampling. Recall that $n_{i,C}$ is the number of observed contacts from RNA $i$ on chromosome $C$. For each *trans*-chromosome $C$, we simply sampled $n_{i,C}$ loci from chromosome $C$ from a multinomial distribution with event probability vector $\{b_j\}$chr($j$)=$C$ .

#### Supplementary Tables

***Table S1.* Differential Expression by total RNA-seq and in the caRNA transcriptome by ChAR-seq, related to Figure 1.** All Gencode V29 genes and UTLs are included, and differential expression was computed for exons, introns, and UTLs separately.

***Table S2.* Chromatin Association scores, related to Figure 1.**

***Table S3.* Catalog of UTLs and their classification, related to Figure 2.**

***Table S4.* Final *trans*- and *cis*-delocalization scores averaged over replicates, related to Figure 4.**

***Table S5.* Catalog of RNAs with extreme delocalization scores, related to Figure 4.**
***Table S6.* List of RNA-DNA contacts not predicted by the generative model, related to Figure 5.** Contacts are defined using 100 kb bins on the DNA side, and individual RNAs (annotated exon, intron, or UTL) on the RNA side. Contacts not predicted by the model were those with a $Log_2$ Fold Change observed over model greater than 1.3 and an adjusted *p*-value less than 0.05, as in Figure 5c,e.

***Table S7.* Coarse graining of Gencode V29 annotations.**

***Table S8.* *trans*-delocalization scores by sample.**

***Table S9.* *cis*-delocalization scores by sample.**
